# Multiplexed construction of defined DNA products from unpartitioned oligonucleotide pools

**DOI:** 10.64898/2026.05.01.722326

**Authors:** Noah Evan Robinson, Jean-Sebastien Paul, Weilin Zhang, Hanqiao Zhang, Sixiang Wang, Arjuna M. Subramanian, Tianhua Zhao, Ezri Abraham, William Ho, Benjamin Simpson, Matt Thomson, Kaihang Wang

## Abstract

Reliable and cost-effective *de novo* DNA production at scale has become increasingly important to studying and engineering biology. Double-stranded DNA is constructed from short synthetic single-stranded DNA oligonucleotides synthesized either individually or as a pool. Oligonucleotide pools offer substantially reduced costs at the sacrifice of yield and individual oligo isolation, making efficient multiplexed DNA construction from pools a complex engineering challenge, yet one which promises to reduce cost, labor, and turnaround time if solved. Here we introduce Oligo Pool Sidewinder assembly, which enables one-pot, parallel assembly of hundreds of DNA fragments simultaneously into dozens of defined constructs with high fidelity from oligo pools. We pair a computational workflow for string-based bespoke oligo design with a set of construction rules for highly multiplexed assembly. We demonstrate recovery of individual sequences from the pool by construct-specific amplification with misconnection rates as low as 1 in 10⁷. This high fidelity for connection enables hundreds of oligos to co-assemble in a single reaction. Further, we show universal amplification of libraries of defined target sequences in a single PCR and extend the approach through *in vitro* hierarchical assembly to construct a large 12.5-kilobase synthetic linear DNA product.

## Main Text

Our ability to design, construct, and test DNA sequences sets the rate at which we study, engineer, and understand biology. Artificial intelligence (AI), large-scale biological datasets, and advanced computational tools have accelerated hypothesis generation to an unprecedented speed and scale, greatly expanding the potential DNA sequence space to explore (*1*). Investigating this diverse sequence space inevitably requires *de novo* DNA construction. However, conventional DNA synthesis and assembly typically handle each sequence individually, resulting in high costs, low throughput, and long turnaround times (*2*). Low-cost, high-throughput, and scalable DNA assembly methods are needed to fully realize the potential of AI-driven biological designs.

Compared with the synthesis of individual oligonucleotides (oligos), the parallel synthesis of multiple oligos in a pool (oligo pools) can substantially increase the scale and reduce the cost of single-stranded DNA (ssDNA) oligonucleotides (*3*). However, these advantages introduce new challenges to DNA assembly such as cross-hybridization between unintended oligo partners, misassembled products, low yield, and an uneven distribution of oligos (*2*). Such challenges can reduce the efficiency and reliability of assembly.

To address these challenges, existing methods for DNA assembly from oligo pools have increased assembly capacity through two complementary approaches: i). partitioning oligo pools into smaller sub-pools, and ii). increasing the number of DNA fragments that can be assembled within a pool. For techniques partitioning oligo pools into smaller sub-pools, large oligo pools can be partitioned through selective amplification (*4*), compartmentalization in droplets (*5*), or optomechanical retrieval of selected oligos (*6*). Creating sub-pools physically isolates assembly reactions into distinct compartments to increase assembly efficiency and reliability. While these approaches decrease assembly complexity, they incur increased cost, workflow complexity, and dependence on specialized equipment (*4–7*).

For techniques increasing the number of DNA fragments that can be assembled within a pool, computational and data-driven design strategies aim to improve assembly reaction accuracy to enable more DNA fragments to be connected orthogonally in parallel (*8, 9 37, 38*). These methods select sets of junction sequences predicted not to cross-hybridize, so that many reactions can proceed independently within a single tube. While these approaches do increase the number of fragments assembled compared to an individual sequence assembly, advancements have largely focused on optimizing Golden Gate assembly (*8, 38*) which relies on 3 or 4 base pair (bp) overhangs that impose a mathematical upper limit of 4^3^ = 64 or 4^4^ = 256 unique overhangs respectively in a single reaction. In practice the usable number is significantly smaller, since not all overhangs in a set are mutually orthogonal (*35*).

The recently developed and highly accurate DNA assembly technique, Sidewinder, has achieved a misconnection rate of nearly 1 in 10^6^ using the DNA 3-Way Junction (3WJ), pointing to a potential to drastically increase the number of fragments assembled simultaneously (*9*). However, the published conventional Sidewinder workflow utilizes only individually synthesized oligos and has limitations that challenge its applicability to oligo pools. These challenges include designing bespoke barcodes with a computationally costly thermodynamics-based algorithm, annealing of individual fragments in isolated tubes, and PAGE purifying individual fragments to remove unannealed oligos. Further, it is not certain whether the low misconnection rate afforded by the conventional Sidewinder reaction will scale with magnitude increases in the number of assembled fragments (from 10s of fragments to 100s).

Here, we introduce Oligo Pool Sidewinder (OPS) assembly, a novel oligo pool-based technique for DNA construction that enhances the specificity of DNA junction interactions, thereby increasing the throughput of oligo pool assembly. OPS achieves this by overcoming the limitations of conventional Sidewinder DNA assembly via a novel bioinformatic and experimental design to enable first-in-class accuracy while assembling large numbers of DNA fragments in oligo pools via the 3WJ principle. This enhanced connection capacity is leveraged for multiplexed parallel one-pot assembly of hundreds of DNA fragments into dozens of defined constructs from a single oligo pool, addressing one of the foundational challenges in oligo pool assembly.

## Results

### OPS fragment design

OPS assembly begins *in silico* with a set of target sequences that are to be constructed. The input target sequences are passed through a bespoke computational workflow, PyWinder, to output the sequence of the component oligos for an OPS assembly (**Fig. 1a-c**). These oligos are then physically synthesized in a unitary oligo pool as input for the subsequent DNA construction via OPS, that outputs physical dsDNA molecules which may be processed for various downstream applications (**Fig. 1a, d**).

**Fig. 1.**
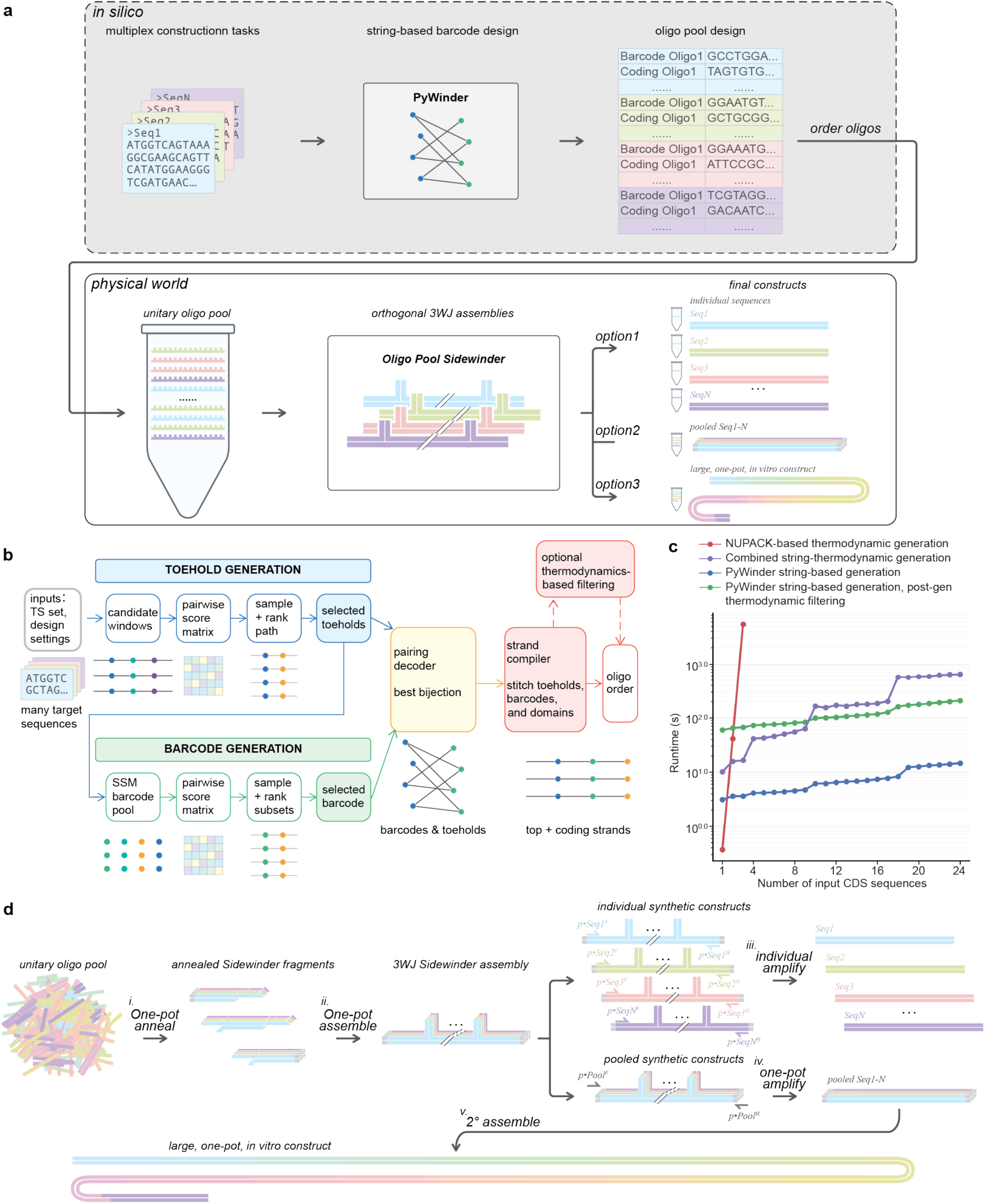
Workflow of Oligo Pool Sidewinder assembly. **a)** Schematic overview depicting the pipeline for Oligo pool Sidewinder (OPS) assembly. Beginning from a set of desired target sequences, PyWinder *in silico* generates the set of oligo sequences that will compose the OPS assembly. The oligos are then synthesized and passed through the OPS protocol to constitute the exact target sequences as physical molecules in the real world. **b)** The PyWinder strand generation workflow. PyWinder takes in target sequences and user-defined settings. Then it identifies a set of mutually orthogonal toeholds and uses these in Sequence Symmetry Minimization (SSM) to generate a set of candidate barcodes that are ranked to identify orthogonal barcode subsets. A pairing decoder then assigns toeholds and barcodes generating a final strand design set. Multiple designs may be optionally generated and undergo thermodynamics-based filtering before a final order-ready design is returned. **c)** Plot comparing the runtime for generating an increasing number of Sidewinder CDS sequences with different Sidewinder barcode design algorithms. **d)** Schematic overview of the OPS protocol. Beginning with oligos synthesized as a unitary oligo pool, all component Sidewinder fragments for all target constructs are annealed in one pot simultaneously (**i**). All 3WJ assemblies are assembled in one pot (**ii**), orthogonally guided by computationally designed barcode sequences. The 2WJ is restored by polymerase activity which can be used in PCR to individually isolate sequences (**iii**), or universally amplify all sequences as a single amplicon (**iv**). This amplicon can then be used for subsequent hierarchical assembly steps for large, *de novo*, *in vitro* constructs (**v**).

All OPS fragments are constituted simultaneously via one-pot annealing (**Fig. 1d, i**). The annealed fragments are subsequently assembled in the one-pot assembly reaction (**Fig. 1d, ii**) into orthogonal 3WJ assemblies, which are directed by highly orthogonal Sidewinder barcode sequences. Then, using either primers specific to each sequence (**Fig. 1d, iii**), or primers universally priming to all sequences (**Fig. 1d, iv**), the 3WJ is removed and target constructs are amplified, either individually in each tube or as a library in one tube, via PCR. These final constructs can then be readily used for any downstream pipeline requiring *de novo* DNA sequences, including being used for one-pot *in vitro* hierarchical construction to larger sizes (**Fig. 1d, v**).

Owing to the increased number of assembly fragments in the same reaction tube, a challenge of OPS is scaling the computational design of orthogonal Sidewinder barcodes. We previously demonstrated assemblies containing 40 fragments connected by 39 barcodes designed using the thermodynamic algorithm NUPACK (*9, 11, 12*). However, scaling the NUPACK multitube design job (*13*) by an order of magnitude became prohibitively computationally expensive (**Fig. 1c, Fig. S1a**). In order to address this challenge, we designed and implemented a string-based Sidewinder strand design workflow, PyWinder, capable of efficiently generating hundreds to thousands of high-performance Sidewinder barcodes in a short period of time (**Fig. 1b, c)**.

At a high level, PyWinder delineates the Sidewinder design procedure as a sequence selection problem on i). candidate toehold loci, ii). barcode sets, and iii). barcode-to-toehold matchings (**Methods**). Our method accepts an arbitrary number of target sequences (TS) as inputs, tiles each TS by overlapping candidate toehold windows and selects one toehold per locus based on a bespoke weighted-Levenshtein edit distance (*14*) metric between possible toeholds (**Methods**), determining the final set of toeholds. PyWinder next designs an extended set of barcodes constrained in sequence against the barcodes themselves and all chosen toehold substrings using Sequence Symmetry Minimization (*15, 16*). From this, PyWinder chooses a subset as the final set of barcodes optimized for conventional Levenshtein edit distance (*14*). Finally, using the final set of toeholds and the final set of barcodes, PyWinder assembles barcode-to-toehold pairings which are selected for shortest Longest Common Substring length in paired toehold-barcode concatenated strings. This generation process creates OPS strands efficiently using purely string-based objectives (i.e. with no thermodynamic criteria) enabling improved speed in a multi-processed manner (**Methods**).

In addition to a purely string-based workflow, we also developed a combined string and thermodynamics-based workflow (**Fig. 1c, Fig. S1b, Methods**). Initial qualitative testing indicated that the faster purely string-based workflow did not perform worse than the combined approach (**Fig. 1c, Fig. S1b, c**). With either method, once multiple complete candidate OPS strand sets were generated, we also developed an optional thermodynamics-based filtering tool to choose the best generated set among an arbitrary number of generations. The optional filter selects over maximum pairwise off target binding probability between different toehold-barcode strands, inspired by previous work leveraging NUPACK (*11, 12, 17*) (**Fig. 1b, c**). The following demonstrations use the purely string-based PyWinder workflow, with endpoint thermodynamic filtering used only where described.

In addition to optimizing for barcode orthogonality, adapting and scaling Sidewinder for OPS requires consideration of homology between target sequences. For conventional Sidewinder, oligos are individually isolated and annealed in separate tubes whereas for OPS, prior to assembly, one-pot annealing is used to direct the formation of all intended OPS fragments simultaneously in a singular pool. This gives rise to the possibility for mis-annealing between unintended oligo partners (**Fig. 2a**) with the potential to decrease the accuracy, yield, and reliability of assembly.

**Fig. 2.**
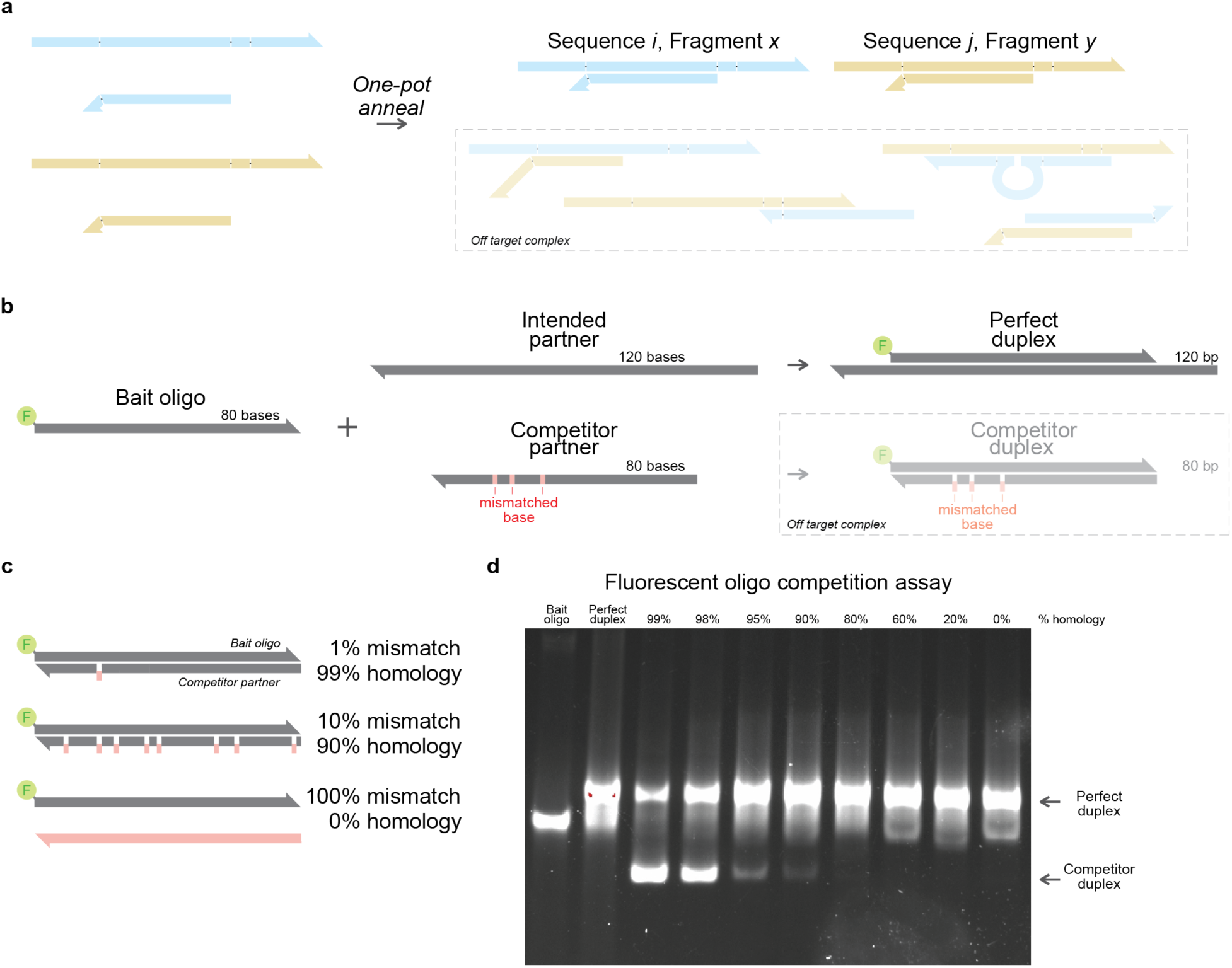
Annealing heteroduplexes in the presence of largely homologous competitors favors the intended partner. **a)** Schematic outlining the challenges presented by one-pot annealing of multiple DNA fragments in the same tube. **b)** A fluorophore tagged bait oligo is annealed in the presence of a perfectly homologous intended partner and an imperfectly homologous competitor partner that contains mismatched bases. Annealing can result in a perfect duplex at 120 bp or a competitor duplex at 80 bp. **c)** A series of competitor oligos are designed with randomly generated mismatched positions. A single mismatch composes the 1% condition, 8 mismatches compose 10% condition and so on through to 100% mismatched which shares no complementarity with the bait oligo. **d)** Bait oligos are annealed with 1:1 concentration of intended partner and 1:10 concentration of competitor partner (intended : competitor) and run on an unstained 8% TBE-PAGE gel and imaged, tracking the migration of fluorophore containing molecules only.

To test the boundary conditions required for mitigating crosstalk between non-partnered oligos, we designed a heteroduplex annealing assay where a fluorophore (Cy3)-tagged bait oligo was annealed in a 1:1 ratio to an intended annealing partner in the presence of a competitor in 10× excess to simulate the type of competition expected in OPS one-pot annealing (**Fig. 2b**). By repeating the assay for a decreasing amount of shared homology between the bait oligo and the competitor, we can estimate the maximum amount of homology that can be shared and tolerated between any two Sidewinder fragments in an oligo pool assembly before substantial mis-annealing of fragments would be observed (**Fig. 2c**). Running each of the annealed conditions on an unstained TBE-PAGE gel allows us to compare annealing by tracking the migration of fluorophore containing molecules only: where an upper band indicates the intended duplex and a lower band indicates the competitor duplex.

Using this assay, we observe that for as high as 95% homology (5% mismatched bases), the intended duplex is the primary product even in the presence of a 10× excess of the competitor (**Fig. 2d**). At 80% homology (20% mismatched bases), we qualitatively observe a minimally detectable presence of the competitor duplex under these conditions. These results are consistent with published literature for similar experiments on heteroduplex annealing (*10*). This indicates that proper formation of the Sidewinder heteroduplex could be highly specific when assembling constructs in a diverse sequence space and that the 80% homology cut off can be used as an empirically validated design rule under the tested conditions.

### Construct-specific amplification of pooled Sidewinder constructs

Once the OPS pooled oligos have been designed *in silico* for orthogonal assembly of an arbitrary number of target sequences via PyWinder, one-pot fragment annealing and one-pot construct assembly is conducted for all sequences simultaneously. To demonstrate this principle, we first constructed 24 target sequences encoding the wildtype coding sequences (CDS) of a set of secreted human proteins (**Fig. 3**). When expressing proteins in their native context (such as human proteins produced in human cell lines) it may be preferable to use the exact native DNA sequence rather than a non-native codon “optimized” sequence (*18, 19*). Owing to Sidewinder’s capacity for constructing uniquely complex DNA sequences (*9*), OPS from oligo pools may offer efficient parallel assembly of potentially complex sequences, including the exact native CDS of human proteins which may contain high GC content, secondary structures, or repetitions.

**Fig. 3.**
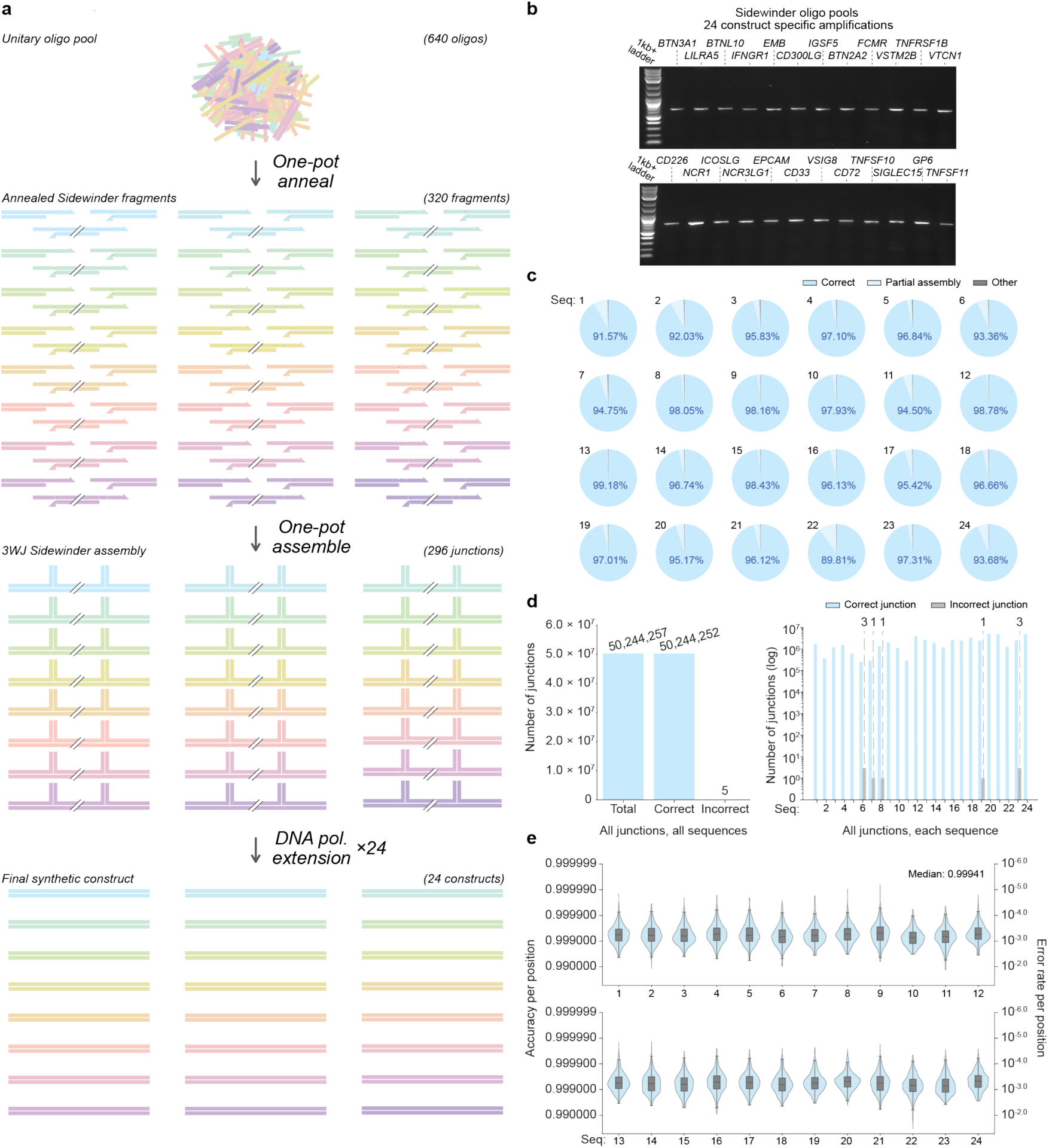
Oligo pools are used to assemble DNA constructs via Sidewinder with high fidelity. **a)** Schematic depicting construct specific amplification of 24 DNA sequences constructed in parallel from oligo pools using Sidewinder. **b)** DNA agarose gel depicting 50 ng of the final PCR products for each of 24 individual constructs with a single strong target band for each. **c)** Fragment-level PacBio sequencing analysis of each of the 24 individual constructs depicting correct assemblies (blue), partially aligned products (a subset of fragments in the correct order) (light blue), and other products composed of PCR artifacts, sequencing artifacts, or incorrect assemblies (grey). Sequence number corresponds to gene name in (**b**) left to right. **d)** Junction-level PacBio sequencing analysis from unfiltered data. Plots depict all observed junctions for all 24 amplicons depicting total junctions (blue), correctly ligated junctions (blue), and incorrectly ligated junctions (grey) with the global count (left) and the count by sequence (right). Inter-construct misconnections (3 between Seq. 6 and Seq. 23, and 1 between Seq. 8 and Seq. 19) are double counted when visualized by individual sequence. One intra-construct misconnection is observed for Seq. 7. We observe a total of 5 misconnections out of 50,244,257 total connections. **e)** Nucleotide-level PacBio sequencing analysis as violin plots and nested box and whisker plots depicting the distribution of per-base accuracies for each position for each of the 24 constructs. Summary statistics available in **Source Data**.

We commercially acquired our unitary oligo pool containing 640 oligos, encoding for 24 orthogonal 13-14 fragment assemblies composing products approximately 650-750 bp in length. All 320 fragments were annealed and assembled in parallel in the same reaction tube (**Fig. 3a**) using thermocycler Assembly protocol 1 (**Methods**). Then using primers that specifically annealed to the coding sequence of each gene, each of the 24 constructs were individually amplified and isolated from the pool in its own construct-specific PCR reaction (**Fig. 3a**). This produced a single, strong, target amplicon of the expected size in all cases with no qualitative sign of mis-assembled byproduct for each of the individually amplified constructs (**Fig. 3b**). This individual construct-specific PCR enables deconvolution from the multiplexed OPS construction into 24 individual constructs with one sequence in each tube.

These results were repeated 2 additional times with slight modifications to the thermocycler protocol (labeled Assembly protocol 2, and 3 respectively in **Methods**) yielding comparable results in all cases. The 24 amplicons for each of the 3 assembly conditions were sent for PacBio sequencing for single molecule high-fidelity reads (*20*). For all assembly conditions, fragment-level analysis shows consistently high rates for correctly assembled constructs (each fragment in the correct order) (**Fig. 3c, Fig. S2**). For Assembly protocol 2, we observe a proportion of correct assemblies ranging from 89.81% correct (sequence 22) to as high as 99.18% correct (sequence 13) (**Fig. 3c**). The major byproduct observed for all constructs was partially assembled sequences (a subset of the assembly fragments with each fragment with the correct connection). Gel extraction of the target bands often reduces the instances of partial reads in the PacBio sequencing data (**Fig. S2a**). The “other” reads are composed of reads not distinguished in type which can be sequencing artifacts, PCR artifacts such as mis-priming during PCR, or potential incorrect ligations at the 3WJ. For the 24 sequences in Assembly protocol 2, “other” reads compose a maximum of 0.41% of reads (sequence 7) (**Fig. 3c**). Assembly protocols 1 and 3 yielded similar results qualitatively on the agarose gel and quantitatively with high global proportions of correct assemblies across all sequences at 97.70% and 95.95% for Assembly protocol 1 and 3 respectively, compared to 96.21% for Assembly protocol 2 and 95.34% for non-gel extracted products with Assembly protocol 1 (**Fig. S2b, c**).

To assess the misconnection rate at the 3WJ more specifically, we applied our junction analysis pipeline to identify all instances of ligated 3WJs that could result from either correct or incorrect ligation (*9*). For 19 of the 24 target sequences with Assembly protocol 2, we observed 0 mis-ligated junctions out of 43,325,799 junctions. The remaining 5 sequences had junction accuracies between 99.9987% and 99.9999%. In total, out of 50,244,257 observed junctions, we observed just 5 mis-ligated junctions (with 4 out of 5 misconnections as inter-sequence mis-ligations) resulting in a post-amplification junction accuracy exceeding 99.99999% or a misconnection rate of 1 in 10,048,851 (**Fig 3d**). Across the 296 junctions for the 24 constructs, we observed an over one order of magnitude lower misconnection rate than was previously reported with the standard Sidewinder protocol and solely thermodynamics-based barcode design algorithm across 9 junctions for a single construct (*9*). The analysis was repeated for the other 3 conditions (Assembly protocol 1, Assembly protocol 1 with no gel extraction, and Assembly protocol 3) which yielded global misconnection rates of 1 in 1,956,782, 1 in 6,572,872, and 1 in 5,837,664, respectively (**Fig. S3)**.

We also characterized the assembly at the nucleotide level, leveraging the per-base accuracy of PacBio sequencing. The median accuracy for a position in any of the 24 target constructs was above 0.999 with a global median of 0.99941, which is approximately 1 mutation for every 1,695 bases (**Fig. 3e, Fig. S4a**). We observed comparable accuracies across each of the other 3 assembly conditions (**Fig. S4a**). Positions intended for adenine (A) and thymine (T) had marginally higher median accuracies than guanine (G) and cytosine (C) for all assembly conditions (**Fig. S4b**). For each of the 24 constructs, we observed a rate of nucleotide perfect clones between 54.149% (sequence 22) and 77.913% (sequence 4). We also observed that, for these oligos from this provider at this target sequence size range, the rate of perfect clones was more strongly correlated to the GC content of the target construct than the length of the target construct (**Fig. S4c, d**). This observation was consistent with each of the replicates for the other 3 assembly conditions (**Fig. S4)**.

These data demonstrate the capacity for OPS to assemble oligos from DNA oligo pools with high instances of correct assembly, low misconnection rates, and high proportions of mutation-free clones.

### Iterative AI protein design and construction via OPS

The rise of AI-informed DNA and protein design has increased the demand for affordable and efficient DNA construction as it is the only way to realize the surplus of *in silico* designed AI sequences in the physical world (*23–27*). The validating, training, and finetuning of biological AI models is an iterative process that can require billions of data points to facilitate machine learning (*30*). Converting the *in silico* DNA and protein designs to physical DNA molecules, which are screened, selected, or sorted based on function, bottlenecks the training and actuation of biological AI-design. The more efficiently designs can be constructed and tested, the more data can be generated to reinforce models for more accurate prediction of sequences which output a desired function.

Improving assembly throughput and readout using OPS helps to facilitate the connection between *in silico* design and empirical functional readout in the physical world (**Fig. 4a**). Towards this end, we redesigned OPS to recover all parallelly assembled target constructs in a single PCR using a single primer pair rather than individually amplifying each target sequence (**Fig. 4b**). Universal amplification can further decrease time and cost for the assembly while increasing throughput. However, reliable universal amplification can only be realized with high fidelity assembly as any misconnections resulting in a truncated product with two terminal fragments may overwhelm the PCR due to PCR’s preference for amplification of shorter products (*9, 21, 22*). The exceedingly low misconnection rate for OPS may afford an improved capacity for robust parallel assembly with universal amplification.

**Fig. 4.**
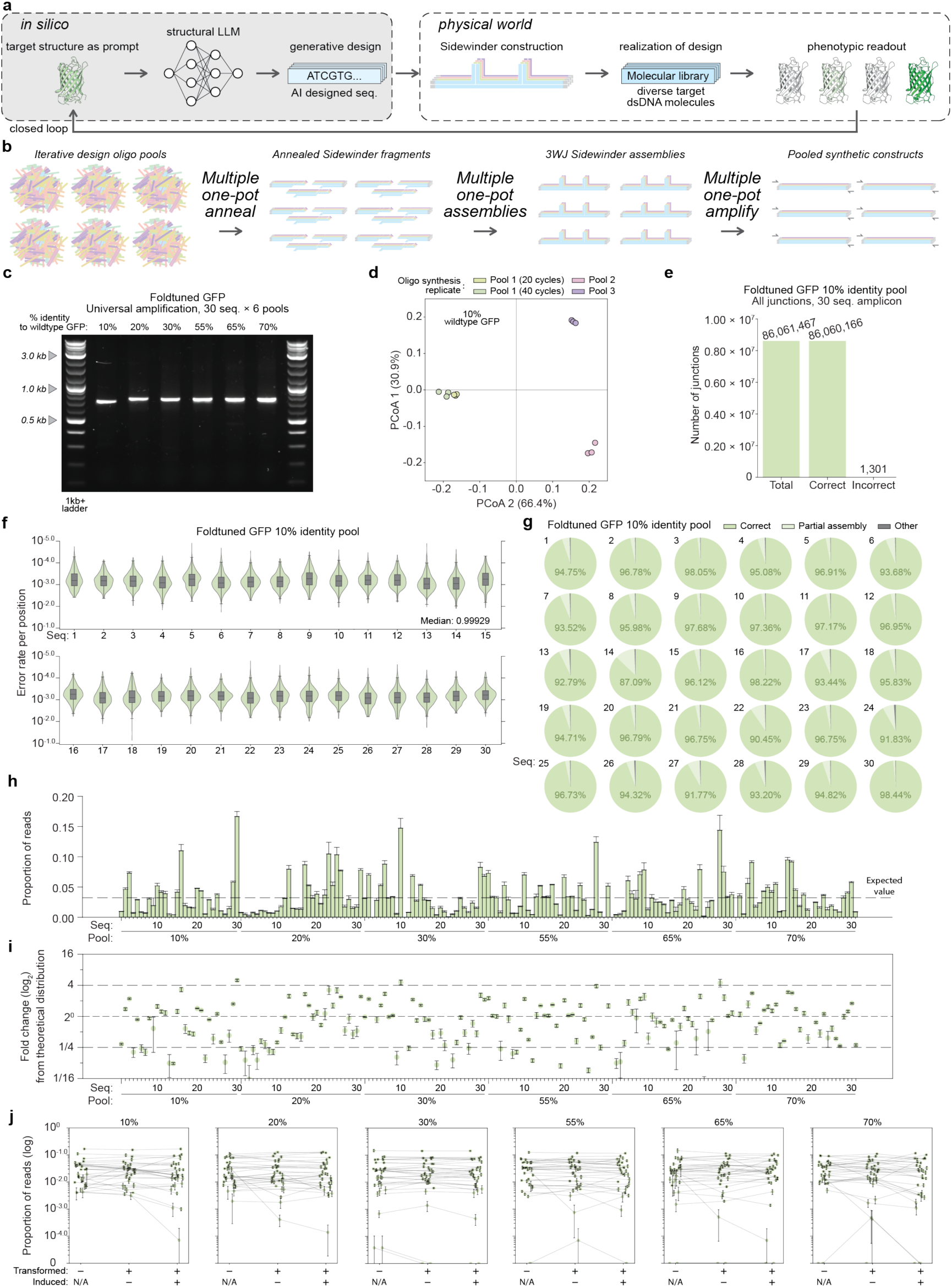
Sidewinder oligo pools enable iterative AI-design of protein sequences in a diverse sequence space. **a)** Schematic depicting Sidewinder facilitating the closed loop pipeline of *in silico* AI designs to physical molecules that may be studied. GFP structure pdb:1ema from Ormö *et al* (*29*). **b)** Schematic depicting universal amplification of DNA sequences constructed in parallel from oligo pools using Sidewinder. **c)** DNA agarose gel depicting 50 ng of the final PCR product after one-pot universal amplification of the 10%, 20%, 30%, 55%, 65%, and 70% wild-type identity constructed pools. **d)** Principal coordinate analysis (PCoA) plot based on Bray–Curtis distances for three replicates of 10% identity 30 sequence design. Pool 1, 2, and 3 are three independently ordered and synthesized oligo pools of the exact same oligo sequences. Each pool was assembled in triplicate. Pool 1 was amplified via PCR with both 20 and 40 cycles. Pools 2 and 3 were amplified with 40 cycles. **e)** Junction-level PacBio sequencing analysis of the 3WJs in the 10% identity universal amplicon depicting the global count with total junctions (green), correctly ligated junctions (green), and incorrectly ligated junctions (grey). **f)** Nucleotide-level PacBio sequencing analysis as violin plots and nested box and whisker plots depicting the distribution of per-base accuracies for each position for each of the 30 constructs in the 10% identity assembly. Summary statistics available in **Source Data**. **g)** Fragment-level PacBio sequencing analysis of the 10% identity universal amplicon divided by construct depicting correct assemblies (green), partially aligned products (a subset of fragments in the correct order) (light green), and other reads composed of either PCR artifacts, sequencing artifacts, or incorrect assemblies (grey). **h)** Bar plot depicts the proportion of Oxford Nanopore sequencing reads assigned to each of the sequences in each of the six AI-designed GFP pools after Sidewinder construction and universal amplification. Each are PCR amplified in triplicate and sequenced. **i)** Dot plot depicts the fold change of the proportion of reads from the expected distribution from that pool (1/30 or 1/31) in the universal amplicon after the 3WJ removal and 40 cycles PCR amplification done in triplicate for each of the six AI-designed GFP pools. Some sequences with low or no representation are not visualized on the plot for legibility but can be found in **Source data**. **j)** Scatter plots depict the proportion of Oxford Nanopore sequencing reads assigned to each of the sequences in each of the six AI-designed GFP pools after Sidewinder construction before cloning, after cloning, and after cloning with induction. A line connects a single target sequence through each step.

Our targets for construction were protein designs generated *in silico* and predicted to fold to a structure similar to GFP within a diverse amino acid sequence space. Because fluorescence reports folding directly, this target lets us test the construction step of an AI-driven design-build-test cycle end to end. We generated 1,530 candidate sequences by foldtuning the ProtGPT2 (31) protein language model using the same methods as in (24). Briefly, foldtuning iteratively i). generated sequences, ii). retained sequences predicted to adopt the target GFP-like fold (TM-score > 0.5), and iii). selected structurally validated sequences for subsequent model updates based on maximal semantic change from natural GFP-like sequences in the embedding space of the ESM2-650M protein language model (*41*), rather than by directly minimizing sequence identity. The resulting *in silico*-designed candidates spanned 5.6-18.5% amino acid sequence identity to wild-type (wt) GFP. For our first round of physical realization, we chose 30 targets from the foldtuned candidates which shared just 10% alignment at the amino acid level to wtGFP but were predicted to fold similarly to GFP in structure space.

We acquired our oligo pool containing 840 oligos, encoding 30 orthogonal 14-fragment assemblies composing products 710 bp in length with a universal priming region at both termini. All 420 fragments were then annealed and assembled in parallel in the same reaction tube and amplified with a single primer pair to produce the universal amplicon with a single, strong, target amplicon of the expected size with no sign of mis-assembly (**Fig. 4c**). As little as 1 fmol of each oligo (20 pM final concentration) is sufficient to yield an amplicon of the expected size (**Fig. S5a**).

Initial investigations showed that only 29 of the 30 sequences were represented in the OPS construction, with sequence #29 absent. To determine the cause of this sequence dropout, we reordered the same 840 oligo pool (Pool 1) two additional times (Pool 2, Pool 3) using the same barcode and coding oligo designs to generate three independent oligo synthesis replicates. We also tested Pool 1 with 40 PCR amplification cycles. The distribution of the 30 parallel-constructed sequences was variable across synthesis replicate (**Fig. 4d**), with one replicate showing as many as 6 dropouts (**Fig. S5b**). The manufacturer-reported oligo dropout rate for their oligo pools is expected to be 0.8% and is reportedly dependent on sequence length and complexity. This would correspond to an estimated 6.7 missing oligos in an 840-oligo pool. One replicate did have representation of all 30 sequences in the universal amplicon (**Fig. S5b, Source Data**).

Performing statistical analysis on the normalized target read proportions in each constructed pool yielded a significant PERMANOVA across all groups (pseudo-*F* = 128.97, p-value = 0.0001), whereas PERMDISP was not significant (*F* = 0.416, p-value = 0.4837) with 9,999 permutations (**Methods**). This indicates that the differences in overall composition across groups are unlikely to be explained primarily by differences in within-group dispersion. Consistently, PCoA based on Bray–Curtis distances showed clear clustering by synthesis replicate with little variation among samples differing only in PCR cycle number (**Fig. 4d**). This suggests that variability during oligo synthesis, including reduced oligo yield or complete oligo dropout, strongly influences the final distribution of targets in the universal amplicon.

Equipped with this understanding, we retried the assembly of Pool 1 supplemented with an equimolar amount of the oligos composing sequence 29. After this supplementation, with all oligos being present in the starting pool, Sidewinder is capable of simultaneously assembling the 420 fragments into each of the corresponding 30 targets with the universal amplicon. For comparison, this successful 390-junction assembly is 1.52× the mathematical limit of the theoretical number of junctions of a single Golden Gate reaction, which is limited by the 256 possible 4-bp overhang combinations (*35*). This further suggests that the fidelity of OPS assembly can exceed the fidelity of some synthesized oligo pools.

Quantifying the assembly at the junction level, per-target junction accuracy across the 30 pooled targets ranged from 99.9432% to 99.9997%, giving a global misconnection rate of 1 in 66,150 (**Fig 4e**). At the nucleotide level, the median accuracy was 0.99930 per position across all constructs with a rate of nucleotide perfect clones between 64.347% (sequence 14) and 77.223% (sequence 5) (**Fig. 4f**, **Fig. S5c**, **Source Data**). At the fragment level, we see consistently high rates for correctly assembled constructs ranging from 87.09% (sequence 14) correct to 98.44% correct (sequence 30), and with a global accuracy of 96.79% correct, 2.89% partial, and 0.31% other reads (**Fig. 4g, Fig. S5d**). None of our initial GFP designs produced a fluorescent phenotype.

Owing to the ease of construction with OPS, we iterated our design process using ProteinMPNN (*32*) to titrate amino acid sequence similarity, producing 5 more sets of 30 sequences each with varying median amino acid sequence alignment to wtGFP-like sequences demarcated into 20%, 30%, 55%, 65%, and 70% identity groups respectively (**Fig. S5f**), plus one verified mScarlet as a positive expression control in each group. Increasing the sequence similarity to wtGFP at the amino acid level also increases the sequence similarity at the DNA level (**Fig. S5g**), demonstrating instances of Sidewinder oligo pools orthogonally assembling similar DNA sequences.

Processed individually, each of the 5 pools, composing a 31-sequence parallel Sidewinder construction with universal amplification, produced a single, strong, target amplicon of the expected size for each (**Fig. 4b, c**). Of the 155 targets, 150 were observed in the Nanopore sequencing (**Fig. 4h**) with 78.7% of constructs being within 4× of the expected distribution (**Fig. 4i**). Transformation of the constructed libraries resulted in consistent representation for each of the 30 sequences within each of the varying percent identity groups before cloning, after cloning, and after cloning with induction (**Fig. 4j**). However, these 150 round two designs likewise did not produce a green fluorescent phenotype.

For the 10% similarity group, the transformation conditions were conducted with the Pool 1 synthesis. Rerunning the PCoA of Bray–Curtis distances for the 10% similarity group showed clustering of both post-transformation conditions with the pre-clonal Pool 1 assembly replicates (**Fig. S5e**). This suggests that for these constructs, oligo synthesis was a stronger determinant of the final distribution of targets in the universal amplicon than transformation and induction.

### OPS and in vitro hierarchical assembly

In addition to producing many variations of smaller sequences to produce a desired function, AI DNA sequence design can produce genome-scale DNA designs *in silico* (23, 26, 36). These larger DNA fragments must also be converted to physical molecules in order to be tested for functionality, similarly bottlenecking genome engineering via AI-design (**Fig. 5a)**. To demonstrate OPS as the technology to meet this demand, we redesigned an essential segment of the *E. coli* MDS42 genome by bidirectionally prompting the Evo 2 model (*23*) with 3 kb segments flanking a 15 kb locus (**Methods**) (**Fig. 5a, b**). Using OPS with universal amplification, our aim was to first produce many multiple defined sequences in parallel in a single PCR with even distribution (**Fig. 5c**). Once this was achieved, we further aimed to conduct a one-pot *in vitro* hierarchical assembly to rapidly construct the AI-designed 12.5 kb DNA molecule (**Fig. 5a-d**).

**Fig. 5.**
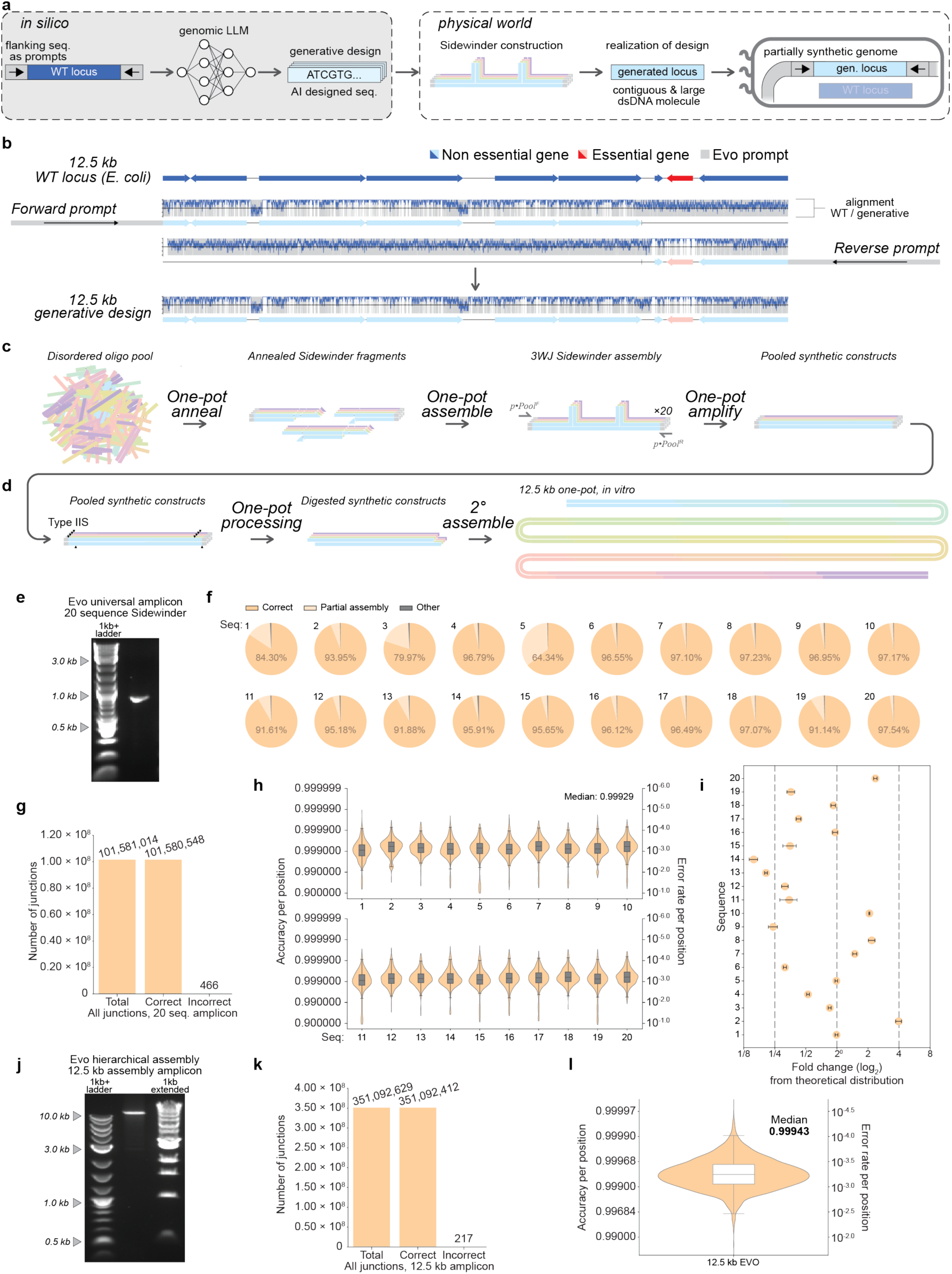
Sidewinder oligo pools have reliable amplification using universal primers, lending to efficient *in vitro* hierarchical construction. Summary statistics available in **Source Data**. **a)** Schematic depicting Sidewinder facilitating the conversion of *in silico* AI designs to physical molecules that may be studied. **b)** Schematic and plot depicting the design of a 12.5 kb segment of the *E. coli* MDS42 genome with non-essential genes (blue) and essential gene (red) to scale. Forward and reverse prompting using Evo 2 are utilized and combined to produce a single 12.5 kb target sequence. Alignment is shown with Smith-Waterman local alignment mapped onto the wildtype reference with a rolling window-smoothed signal. **c)** Schematic depicting universal amplification of DNA sequences constructed in parallel from oligo pools using Sidewinder. **d)** Schematic depicting the single amplicon, containing a library of defined sequences, assembled in a subsequent round of hierarchical assembly into a large, linear dsDNA product. **e)** DNA agarose gel depicting 50 ng of the final PCR product after one-pot universal amplification of the constructed pool, containing each of 20 target constructs in a single strong target band. **f)** Fragment-level PacBio sequencing analysis of the universal amplicon divided by construct depicting correct assemblies (orange), partially aligned products (a subset of fragments in the correct order) (light orange), and other reads composed of either PCR artifacts, sequencing artifacts, or incorrect assemblies (grey). **g)** Junction-level PacBio sequencing analysis of the 3WJs in the universal amplicon depicting the global count with total junctions (orange), correctly ligated junctions (orange), and incorrectly ligated junctions (grey). **h)** Nucleotide-level PacBio sequencing analysis as violin plots and nested box and whisker plots depicting the distribution of per-base accuracies for each position for each of the 20 constructs. **i)** Dot plot depicts the proportion of Oxford Nanopore sequencing reads assigned to each of the 20 sequences divided by the theoretical proportion (1/20 = 0.05) in the universal amplicon after the 3WJ removal and 20 cycles PCR amplification done in triplicate. **j)** DNA agarose gel depicting 50 ng of the final PCR product for the 12.5 kb assembly amplicon after *in vitro* hierarchical assembly with a single strong target band. **k)** Junction-level PacBio sequencing analysis of the 3WJ after construction to 12.5 kb depicting the global count with total junctions (orange), correctly ligated junctions (orange), and incorrectly ligated junctions (grey). **l)** Nucleotide-level PacBio sequencing analysis as a violin plot and nested box and whisker plot depicting the distribution of per-base accuracies for each position of the 12.5 kb assembly amplicon for only reads aligning to the full 12.5 kb sequence.

We acquired our oligo pool containing 600 oligos, encoding for 20 orthogonal 15-fragment assemblies composing products 728 bp in length. The targets were designed with a shared priming region at both termini external to a non-coding buffer region ensuring all constructs were exactly 728 bases in length to avoid amplification bias due to PCR (**Fig. S6a**). All 300 fragments were then annealed and assembled in parallel in the same reaction tube and amplified with a single primer pair to produce the universal amplicon containing all 20 constructed sequences. This process produced a single, strong, PCR product of the expected size with no sign of mis-assembly (**Fig. 5e**).

Investigating this product with PacBio sequencing shows that each of the 20 targets are represented in the universal amplicon, demonstrating successful construction and pooled amplification using Sidewinder from oligo pools. Quantifying the assembly at the fragment level shows high rates of correctly assembled constructs up to 97.54% correct (sequence 20) with a global accuracy of 91.28% correct, 8.25% partial, and 0.48% other reads (**Fig. 5f, Fig. S6b**). At the junction level, we observe a junction accuracy of 99.9995% or a misconnection rate of 1 in 217,985 (**Fig. 5g**). The increase in 3WJ misconnection was observed to be higher for both universal amplification conditions which may be attributed to the profile of misconnections that may be amplified with universal amplification and the large number of assembly junctions (**Fig. S7**). Lastly, at the nucleotide level, the median accuracy was 0.99929 per position across all 20 constructs with a rate of nucleotide perfect clones between 64.616% (sequence 11) and 76.490% (sequence 18) (**Fig. 5h, Fig. S6c**).

We also assessed the distribution of the proportion of reads assigned to each of the 20 sequences to quantify the composition of the pooled assembly. Variation in the distribution may result from multiple sources including differences in yield during oligo synthesis, PCR bias during amplification, sequencing bias during sequencing, in addition to assembly bias during construction. Given an expected proportion of 1/20 = 0.050 for each of the 20 targets, the least represented sequence (sequence 14) was approximately 6× underrepresented with an average proportional occurrence of 0.008 and the most represented sequence (sequence 2) was approximately 4× overrepresented with an average proportional occurrence of 0.198 (**Fig. 5i**). Changing the number of PCR cycles did not influence the shape of the distribution (**Fig. S6d**).

The quality, purity, and relatively even representation of each target construct in the universal amplicon enables the amplicon to be further processed for hierarchical assembly entirely *in vitro* (**Fig. 5d**). Golden Gate was used for the higher order assembly to facilitate scarless removal of the universal priming and buffer regions using a Type IIS enzyme that cuts internally to expose short 4 bp overhangs on each of the 20 constructs (**Fig. S6a**). Two rounds of *in vitro* hierarchical assembly with a final amplification of the assembly yields a single strong target band of the correct size (**Fig. 5j**).

We applied our assembly characterizations to the 12.5 kb *in vitro* hierarchically assembled product to compare the influence the hierarchical assembly had on the constructed molecule. When conducting the exact same junction analysis as at the universal amplicon level, the Sidewinder 3WJ misconnection rate was observed to be 1 in 1,617,938 (**Fig. 5k**). At the nucleotide level, we measured a median per-base accuracy of 0.99943 considering only reads with >80% alignment to the full 12.5 kb amplicon (**Fig. 5l**) whereas we previously measured the median per-base accuracy as 0.99929 for the twenty 728-base constructs in the universal amplicon (**Fig. 5h**). Comparing the mutation frequencies indicates that hierarchical assembly does not increase the mutation frequency, once again suggesting that most SNPs in the final construct likely originate from oligonucleotide synthesis. In total, despite the large size and the entirely *in vitro* construction, we identified 96 reads which were mutation-free in the PacBio sequencing dataset.

## Discussion

Here, we introduce Oligo Pool Sidewinder assembly which adapts the 3WJ-mediated Sidewinder DNA assembly technology to oligo pools in order to simultaneously connect hundreds of DNA fragments into defined constructs in a single tube from a single oligo pool. We reproducibly construct target sequences in parallel with high fidelity and representation using both specific amplification for individual construct isolation and universal amplification of all constructs in one pot, producing defined DNA sequences compatible with downstream pipelines for characterization. The demonstrations here show a streamlined protocol for orthogonal sequence construction from a single pool in a single compartment, but OPS is in principle compatible with any upstream oligo pool compartmentalization technology which may allow for synergistic protocols that synthesize orders of magnitude more constructs than any individual technique alone.

Utilizing our novel OPS assembly paired with a novel efficient string-based bespoke strand design workflow, PyWinder, we have demonstrated instances of misconnection at the 3WJ to be as low as 1 in 10,048,851 when using construct-specific amplification. This exceedingly low misconnection rate enabled over an order of magnitude more constructs to be made in parallel compared to Sidewinder from individual oligos (*9*) such that in a single reaction, we observed a 10× reduction in measured misconnection rate while achieving over 10× the number of DNA fragments assembled simultaneously. We hypothesize that the lower misconnection rate observed in PyWinder string designs compared to previous purely thermodynamic designs may be due to thermodynamic criteria describing equilibrium ensembles directly whereas the 3WJ formation during Sidewinder is at least partially kinetically controlled. Ultimately, the fidelity of the 3WJ design, the expansive sequence space available for barcode design, and the ease of OPS assembly point to the potential for further scaling to hundreds or even thousands of constructs assembled simultaneously in one-pot from oligo pools. At that scale, sequence accuracy is now likely to be primarily limited by oligo synthesis fidelity rather than by the assembly method itself.

Enabled by the enhanced capacity for design from PyWinder and the enhanced capacity for construction with OPS, we successfully and reproducibly constructed the native wildtype sequences of 24 human genes in a one-pot reaction with construct-specific isolation via PCR amplification. This capacity can be applied to various contexts but is particularly useful when individual nucleotide-perfect sequences are desired. This may facilitate a future of rapid and affordable DNA construction where sequences need not be limited by complexity. In this future, codon optimization becomes a deliberate choice for heterologous expression rather than a workaround for the difficulty of synthesizing native sequences.

In addition to construct-specific PCR amplification, we also successfully constructed dozens of defined sequences in parallel with universal amplification, achieving high representation of constructs and a distribution within 4× the theoretical yield in most cases. This capacity can be applied in a variety of contexts but is particularly useful for downstream pipelines that involve high-throughput readout of many clones with sorting or selection without the need for individual isolation of each construct. We anticipate that OPS will be important for the future of AI-facilitated design as it enables fast, scalable, and robust realization of biological AI designs.

In particular, we showed how this technology can facilitate iterative design of AI-generated sequences. Through screening over 180 constructs in a diverse sequence space that are closely related in structure space, we constructed genes encoding protein products predicted to fold to structures of fluorescent proteins with as little as 8.4% and as high as 71.1% sequence similarity to wtGFP. This capacity will help accelerate our understanding of the connections between sequence, structure, and function by enabling the generation and characterization of novel proteins designed by biological AI models. This work characterizes OPS as a part of an envisioned design-construct-readout-feedback cycle for DNA and protein engineering as notably, none of the 180 designs realized by OPS into physical constructs yielded functional fluorescence when transformed into *E. coli* cells despite being predicted *in silico* to well mimic the structure of GFP. This may be expected as there are known difficulties with predictive folding of GFP-like structures (*39, 40*), showcasing the limitations of *in silico* validation in biomolecular design today, and highlighting the importance of design actualization to further improve generative models.

Further, we showed how OPS enables DNA construction of an *in silico* AI design from 600 oligos in a unitary oligo pool to a final size of 12.5 kb with a novel protocol for *in vitro* hierarchical assembly. Such rapid construction of large DNA from oligo pools offers a potential path forward for the efficient *de novo* synthesis of genomes. Additionally, hierarchical assembly to smaller sizes of 1-10 kb can be implemented for library generation of larger genes beyond the current limits of pooled DNA assembly (*8, 28*).

While the yield and accuracy of construction demonstrated in this study points to the potential for further scaling of the number of sequences assembled in parallel, more investigation must be done into the source of the increased misconnection rate observed with universal amplification. Regardless of the technique used in assembly, universal amplification has the potential to amplify more types of misassembled junctions. Construct-specific amplification is more stringent: a misconnection survives only if it retains both the first and last fragment of the specific targeted construct, since those are what the primers bind to. Universal primers do not impose this same requirement, so a broader range of misassembled products can be amplified, including just a single misconnection between any two constructs. This hypothesis is supported by the resultant decrease in misconnection rate back to under 1 in 10^6^ when the 20-sequence universal amplicon was used for *in vitro* hierarchical assembly that contained a specific primer amplification step.

These results suggest that OPS has the potential to be an important tool in the synthetic biology and bioengineering toolbox as the technique can be interfaced with other pooled construction and genetic engineering methods to better study and engineer biology. We envision that OPS will enable the cost-effective and accessible construction at scale of large DNA libraries, with the potential for further scaling through oligo pool partitioning. Such capacity would rapidly expand today’s testable sequence space and drive advances across agriculture, medicine, materials science, data storage, and beyond.

## Author contributions

Conceived the experimental design and applications: KW and NER

Developed PyWinder: JSP with input from NER and KW

Performed DNA assembly experiments: NER, WZ, TZ, EA and WH

Developed informatic pipelines for the analysis of sequencing data: HZ with input from NER and KW

Developed the foldtuning model and designed AI-sourced GFP: AMS and MT

Curated targets for construction: WZ, BS and NER

Wrote the manuscript: NER, JSP, SW, and KW with input from all of the authors.

We thank W. M. Wojtowicz and K. Zinn for insightful discussion and providing the 24 secreted proteins as targets for construction. K.W. acknowledges funding support from the Caltech Center for Environmental Microbial Interactions (CEMI).

## Data, and Materials availability

The sequencing data generated in this study have been deposited in the NCBI Sequence Read Archive with the submission number PRJNA1491617.

Source data are provided for this paper.

Supplementary information is available for this paper. Correspondence and requests for materials should be addressed to.

## Supplementary Materials

Materials and Methods

References (33–34)

Figs. S1 to S7

Tables S1 to S2

Source Data

## References

1. Koh, H.Y., et al. “AI-Driven Protein Design.” Nature Reviews Bioengineering, vol. 3, no. 12, Dec. 2025, pp. 1034–56. 10.1038/s44222-025-00349-8.

2. Hoose, A., et al. “DNA Synthesis Technologies to Close the Gene Writing Gap.” Nature Reviews Chemistry, vol. 7, no. 3, Jan. 2023, pp. 144–61. DOI.org (Crossref), 10.1038/s41570-022-00456-9.

3. Kosuri, S., and Church, G.M. “Large-Scale de Novo DNA Synthesis: Technologies and Applications.” Nature Methods, vol. 11, no. 5, May 2014, pp. 499–507. DOI.org (Crossref), 10.1038/nmeth.2918.

4. Kosuri, S., et al. “Scalable Gene Synthesis by Selective Amplification of DNA Pools from High-Fidelity Microchips.” Nature Biotechnology, vol. 28, no. 12, Dec. 2010, pp. 1295–99. DOI.org (Crossref), 10.1038/nbt.1716.

5. Plesa, C., et al. “Multiplexed Gene Synthesis in Emulsions for Exploring Protein Functional Landscapes.” Science, vol. 359, no. 6373, Jan. 2018, pp. 343–47. DOI.org (Crossref), 10.1126/science.aao5167.

6. Cho, N., et al. “High-Throughput Construction of Multiple Cas9 Gene Variants via Assembly of High-Depth Tiled and Sequence-Verified Oligonucleotides.” Nucleic Acids Research, vol. 46, no. 9, May 2018, pp. e55–e55. DOI.org (Crossref), 10.1093/nar/gky112.

7. Wan, W., et al. “High-Fidelity de Novo Synthesis of Pathways Using Microchip-Synthesized Oligonucleotides and General Molecular Biology Equipment.” Scientific Reports, vol. 7, no. 1, July 2017, p. 6119. DOI.org (Crossref), 10.1038/s41598-017-06428-0.

8. Freschlin, C.R., et al. “Scalable and cost-efficient custom gene library assembly from oligopools.” Sci. Adv. 12, eady2279 (2026).

9. Robinson, N.E., Zhang, W., Ghosh, R., et al. “Construction of complex and diverse DNA sequences using DNA three-way junctions.” Nature 651, 491–500 (2026).

10. Koehler, R. T., and N. Peyret. “Thermodynamic Properties of DNA Sequences: Characteristic Values for the Human Genome.” Bioinformatics, 21(16), 3333–39 (2005).

11. Fornace, M. E. et al. “NUPACK: analysis and design of nucleic acid structures, devices, and systems.” Preprint at 10.26434/chemrxiv-2022-xv98l (2022).

12. Zadeh, J. N. et al. “NUPACK: analysis and design of nucleic acid systems.” J. Comput. Chem. 32, 170–173 (2011).

13. Wolfe, B. R., Porubsky, N. J. Zadeh, J. N., et al. “Constrained multistate sequence design for nucleic acid reaction pathway engineering.” J Am Chem Soc, 139,3134-3144, (2017).

14. Levenshtein, V. I. “Binary codes capable of correcting deletions, insertions, and reversals.” Sov. Phys. Dokl. 10, 707–710 (1966).

15. Gowri, G., Sheng, K. & Yin, P. “Scalable design of orthogonal DNA barcode libraries.” Nat Comput Sci 4, 423–428 (2024).

16. Dirks, R. M. “Paradigms for computational nucleic acid design.” Nucleic Acids Res. 32, 1392–1403 (2004).

17. Paul, J.S., et al. “Ortho: A multiprocessed orthogonal DNA library generator.” GitHub. https://github.com/jspaul2003/ortho (2024).

18. Gustafsson, C., Govindarajan, S., Minshull, J. “Codon bias and heterologous gene expression.” Trends Biotechnol. 22, 346–353 (2004).

19. Sauna, Z., Kimchi-Sarfaty, C. “Understanding the contribution of synonymous mutations to human disease.” Nat Rev Genet 12, 683–691 (2011).

20. Marx, V. “Method of the year: long-read sequencing.” Nat. Methods 20, 6–11 (2023).

21. Dabney, J. & Meyer, M. “Length and GS-Biases during sequencing library amplification: a comparison of various polymerase-buffer systems with Ancient and modern DNA sequencing libraries.” Biotechniques 52, 87–94 (2012).

22. Hsiau T.H.C., et al. “A Method for Multiplex Gene Synthesis Employing Error Correction Based on Expression.” PLOS ONE 10(3), (2015).

23. Brixi, G., Durrant, M.G., Ku, J., et al. “Genome modelling and design across all domains of life with Evo 2.” Nature (2026).

24. Subramanian, A. M., et al. “Unexplored regions of the protein sequence-structure map revealed at scale by a library of foldtuned language models.” Preprint at bioRxiv 10.1101/2023.12.22.573145 (2025).

25. Shanker, V. R. et al. “Unsupervised evolution of protein and antibody complexes with a structure-informed language model.” Science 385, 46–53 (2024).

26. Nguyen, E., et al. “Sequence modeling and design from molecular to genome scale with Evo.” Science 386, eado9336 (2024).

27. Wolfsberg, E. et al. “Machine-guided dual-objective protein engineering for deimmunization and therapeutic functions.” Cell Syst. 16, 101299 (2025).

28. Zou, S., Wu, Z. & Xu, C. “Hierarchical assembly of long DNA libraries from short oligonucleotide pools.” ICLR 2025 Workshop on Machine Learning for Genomics Explorations. (2025).

29. Ormö, M., et al. “Crystal Structure of the *Aequorea victoria* Green Fluorescent Protein.” Science 273, 1392–1395 (1996).

30. Nijkamp, E., et al. “ProGen2: Exploring the boundaries of protein language models” Cell Sys, 14(3), 968–978 (2023).

31. Ferruz, N., Schmidt, S. & Höcker, B. “ProtGPT2 is a deep unsupervised language model for protein design”. Nat Commun 13, 4348 (2022).

32. Dauparas, J., et al. “Robust deep learning–based protein sequence design using ProteinMPNN”. Science. 378, 49–56 (2022).

33. Daily, J. Parasail: SIMD C library for global, semi-global, and local pairwise sequence alignments. BMC Bioinformatics, 17(1), 1–11 (2016) doi:10.1186/s12859-016-0930-z

34. Altschul, S. F., Gish, W., Miller, W., Myers, E. W., & Lipman, D. J. Basic local alignment search tool. Journal of Molecular Biology, 215(3), 403–410 (1990).

35. Potapov, I., et al. “Comprehensive Profiling of Four Base Overhang Ligation Fidelity by T4 DNA Ligase and Application to DNA Assembly”. ACS Synth. Biol. 7(11), 2665–2674 (2018).

36. King, S. H., et al. “Generative design of bacteriophages with genome language models”. Science. 393, eaec2657 (2026).

37. Sidore, M. A., et al “DropSynth 2.0: high-fidelity multiplexed gene synthesis in emulsions”. Nucleic Acids Research. 48(16), e95 (2020).

38. Romanowicz, K. J., Hilton, S. R., Villegas, N., & Plesa, C. “DropSynth-Gold: Golden Gate Assembly in Emulsions Extends Multiplexed Gene Libraries to Greater Lengths”. bioRxiv (2026).

39. Nam, K. H. “Evaluation of AlphaFold3 prediction for post-translational modification, oligomeric assembly, and quenchable metal binding of fluorescent proteins”. J. Mol. Graph. Model. 142 (2026).

40. Devkota, K., Shonai, D., Mao, J. et al. Miniaturizing and modifying natural proteins with Raygun. Nature (2026).

41. Zeming Lin et al. Evolutionary-scale prediction of atomic-level protein structure with a language model. Science 379, 1123–1130 (2023).

