## Supplementary Materials for "Multiplexed construction of defined DNA products from unpartitioned oligonucleotide pools"

### Materials and Methods

#### String-based Sidewinder strand generation

Using 5' to 3' notation, our string-generator algorithm takes as input a set of target sequences (TSs) $\mathcal{S} = \{s^{(1)}, \ldots, s^{(M)}\}$, and the variables: target oligo length $L$, barcode length $b$, toehold length $t$, and toehold search range $R$. Given this input, the algorithm aims to break down each TS into a set of domains, barcodes, and toeholds which will compose oligos compatible with pooled Sidewinder construction that, when assembled, exactly reconstruct each TS. Chosen toeholds, barcodes, and toehold-barcode concatenations must be 'orthogonal' as defined by satisfying certain string-based criteria (defined below). Thus, the algorithm is decomposed into 3 subprocesses: (1) toehold-set selection, (2) barcode-set selection, and (3) toehold-barcode matching selection.

Toehold-set selection is done on a TS-by-TS basis. Given $b, t, R$, we first define fragment length $f = L - 2b$ and maximum C-domain (paired bases between barcode oligo and coding oligo) length $c_{\text{max}} = L - b$. Next, the first part of toehold-set selection involves generating multiple loci from each TS where each locus contributes exactly $R$ candidate toeholds and downstream selection picks exactly one candidate from each locus. For a TS sequence $s$ of length $|s|$, we initialize a starting locus position (defined as locus 0) $p_0 = L - b - t$. In general, for any locus $m$, we collect the following set of candidate toeholds: $T_m = \{s[p_m + r : p_m + r + t] \;:\; r = 0, 1, \ldots, R - 1\}$. A subsequent locus $m + 1$ can be defined using starting position $p_{m+1} = p_m + f$, unless the break condition $|s| - p_{m+1} \leq c_{\text{max}}$ is satisfied (i.e. once the remaining trailing domain would no longer be longer than $c_{\text{max}}$ so we stop looking for new toehold sets).

Several constraints on the input parameters are implied by our toehold enumeration procedure. First, to ensure that the very first toehold window starts inside the TS, we require $p_0 = L - b - t \geq 0$ (using 0-indexing over strings), which gives $L \geq b + t$. Second, because adjacent candidate groups are separated by $f = L - 2b$, non-contiguous overlap of two candidate windows from loci $m$ and $m+1$ requires $p_m + f > p_m + (R-1) + t$, or equivalently $L - 2b > R + t - 1$. Hence we have a safe integer condition $L > 2b + t + R - 1$. Observe also that the toehold window searching nature of our algorithm dictates the length of the final generated Sidewinder oligos which fall $\in [L-R+1, L-R+2, ..., L+R-1]$. In our work, unless otherwise defined, we used $(b, t, R) = (22, 10, 15)$, which gives $L > 68$. We thus used a safe $L = 96$, giving a min and max final oligo length of $82$ and $110$ nucleotides respectively.

In toehold selection, the string objective computes a weighted edit-distance matrix across all candidate toeholds from different loci. This metric is calculated only between toeholds from different loci. The underlying dynamic program is Levenshtein (*14*)-like, but the penalty cost of an edit at normalized sequence position $u = i/(t-1)$ (with regular sequence position in the toehold $i \in [0, 1, ..., t-1]$) is weighted by $w(u) = 1 + e^{-u}$, so that mismatches at positions closer to 0 (the site of ligation) carry larger distance. This is done to penalize in particular off-target binding at the Sidewinder fragment-toehold junction which has a higher probability of yielding misincorporation from a misligation event. In this way, we use the weighted Levenshtein Edit Distance metric as a model for a worst-case binding scenario when aligning off-target toeholds. Importantly, because this weighted distance operator is inherently non-commutative ($d_w(x, y) \neq d_w(y, x)$), two sequences $x$ and $y$ have their weighted edit distance matrix entries ($d_{toe}(x, y), d_{toe}(y, x)$) symmetrized such that $d_{toe}(x, y) = d_{toe}(y, x) = \min\{d_w(x, y), d_w(y, x)\}$.

Observe that since we only take one toehold from each locus, we only compute cross-locus comparisons. Thus if there are $G$ loci and $R$ candidates per locus, the number of scored toehold edges is $\binom{GR}{2} - G\binom{R}{2}$. Each complete toehold solution is a path containing one candidate from every locus. The algorithm samples $K$ such paths ($K = 1000$ by default in our *in silico* generations). Paths are generated locus-by-locus with the locus considered at each iteration picked randomly. For the first locus, a toehold is picked uniformly randomly, however for future iterations, if the current partial path contains vertices $\{v_1, v_2, \ldots, v_z\}$ and the next locus to-be-sampled-from contains candidates $\{u_1, u_2, \ldots, u_R\}$, then the next toehold to be picked $u_j$ will be chosen via a weighted random sampling scheme. The sampling weight of candidate $u_j$ is $\omega(u_j) = 1 + \sum_{\ell=1}^{z} \exp(d_{toe}(v_\ell, u_j))$. Thus large edit distances are rewarded through the $\exp(d)$ term such that candidates more dissimilar to the existing path are more likely to be sampled. For each completed sampled path $P$, the string score is the tuple $S_{toe}(P) = (\min_{i<j} d_{toe}(P_i, P_j), \frac{1}{\binom{|P|}{2}} \sum_{i<j} d_{toe}(P_i, P_j))$. The final toehold path is chosen by weighted rank aggregation, with default weights 1.0 on the minimum edit distance and 0.5 on the average edit distance; both metrics are maximized (the first and second components of the $S_{toe}$ tuple respectively). We chose 1.0 and 0.5 as our ranking weights in order to both acknowledge the importance of minimum and average edit distance but to also prioritize the minimum since a single misligation event has the potential to greatly disrupt the required orthogonality of the Sidewinder assembly reaction.

Given the selected toehold set $\mathcal{T}$, we next generate an initial barcode pool using a Sequence Symmetry Minimization (SSM) constraint. SSM can be used to generate a set of DNA sequences of length $l$ which do not share a common substring of length $k$. This can be expanded to include reverse complements and shared common substrings of length $q$ with another set of DNA sequences (including that set's reverse complements as well). In our case, the default automatic forbidden substring length between barcodes and $\mathcal{T}$ is $q = \lfloor t/2 \rfloor$ and within the barcodes is $k = \max(\lfloor b/4 \rfloor, q+1)$. Every reverse complement is considered also. To do this, we use seqwalk (*15*), a novel and efficient method to generate maximally sized SSM-constrained DNA sequences.

Our program requires a pool of at least $5|\mathcal{T}|$ admissible barcodes. If seqwalk returns fewer than this number of barcodes under the default $(q, k)$ choice, we relax the SSM constraint and re-run, iterating until the pool is large enough. Indexing such iterations from $i = 0$, we update $(q, k)$ between runs by alternating: on even iterations or when $q \geq t$ we increment $k \leftarrow k + 1$ (admitting longer shared substrings between barcodes); on odd iterations while $q < t$, we increment $q \leftarrow q + 1$ (admitting longer shared substrings between barcodes and $\mathcal{T}$). This alternation tends to keep $k$ and $q$ balanced so that neither the intra-barcode nor the barcode–toehold constraint is relaxed disproportionately. Note that this does mean it is possible, for sufficiently large $\mathcal{T}$ and sufficiently small $b$ that it may not be possible to satisfy our $5|\mathcal{T}|$ criterion, even if we reach an undesirable point where $q = t$ and $k = b$, in which case the program halts unless that $5|\mathcal{T}|$ threshold is lowered.

Barcode-barcode dissimilarity is scored using conventional (unweighted) Levenshtein edit distance (*14*) (represented here by $d_{bar}$). Each barcode is treated as its own singleton group, so this stage effectively samples barcode subsets of size $n = |\mathcal{T}|$ and ranks these subsets. If a sampled subset is denoted $B = \{b_1, \ldots, b_n\}$, its score is represented by the tuple $S_{bar}(B) = (\min_{i<j} d_{bar}(b_i, b_j), \frac{1}{\binom{n}{2}} \sum_{i<j} d_{bar}(b_i, b_j))$, followed by the same weighted rank aggregation used for toeholds' weighted edit distance-based scores. Let $\mathcal{B}$ represent the final picked set of barcodes.

Once the final toehold set $\mathcal{T}$ and the final barcode set $\mathcal{B}$ have been chosen, the algorithm samples $K = 1000$ random bijections between the two. For any bijection with some bijective map $\pi : [1, 2, \ldots, n] \to [1, 2, \ldots, n]$ and defining $\tau_i$ as the $i$th toehold in $\mathcal{T}$, and $\beta_i$ as the $i$th barcode in $\mathcal{B}$, we induce a list of concatenated toehold-barcode sequences $z_i = \tau_i \bullet \beta_{\pi(i)}, \forall i \in [1, 2, \ldots, n]$. The score of that bijective mapping is simply represented by the length of the longest common substring (LCS) found among all pairs $\{z_i\}$. We pick the mapping which minimizes this LCS (or randomly among the mappings which have an equally lowest LCS).

After the global toehold set, barcode set, and toehold–barcode matching have been selected, the chosen pairs are partitioned back according to the TS from which each toehold originated, and the resultant Sidewinder oligos are composed separately for each TS. For a given TS $s^{(r)}$, suppose the selected toeholds assigned to that TS occur at positions $p_1^{(r)} < \cdots < p_{n_r}^{(r)}$, with corresponding matched toehold-barcode pairs $(\tau_1^{(r)}, \beta_1^{(r)}), \ldots, (\tau_{n_r}^{(r)}, \beta_{n_r}^{(r)})$. These positions partition $s^{(r)}$ into $n_r + 1$ inter-toehold C-domains: $D_1^{(r)} = s^{(r)}[0 : p_1^{(r)}]$, $D_i^{(r)} = s^{(r)}[p_{i-1}^{(r)} + t : p_i^{(r)}]$ for $2 \leq i \leq n_r$, and $D_{n_r+1}^{(r)} = s^{(r)}[p_{n_r}^{(r)} + t : |s^{(r)}|]$. The program then constructs $n_r + 1$ Sidewinder barcode and coding strands for that TS. The first barcode strand is barcode$_1^{(r)} = D_1^{(r)} \bullet \tau_1^{(r)} \bullet \beta_1^{(r)}$, while for $2 \leq i \leq n_r$, barcode$_i^{(r)} = \widetilde{\beta_{i-1}^{(r)}} \bullet D_i^{(r)} \bullet \tau_i^{(r)} \bullet \beta_i^{(r)}$, and the final top strand is barcode$_{n_r+1}^{(r)} = \widetilde{\beta_{n_r}^{(r)}} \bullet D_{n_r+1}^{(r)}$ where $\widetilde{\cdots}$ denotes reverse complement. The corresponding coding strands are coding$_1^{(r)} = \widetilde{D_1^{(r)}}$ and for $2 \leq i \leq n_r + 1$, coding$_i^{(r)} = \widetilde{D_i^{(r)}} \bullet \widetilde{\tau_{i-1}^{(r)}}$.

### Thermodynamic Sidewinder strand generation

Thermodynamic strand generation preserves the same TS partitioning, toehold-window enumeration, Sequence Symmetry Minimization barcode generation, and final strand-construction rules defined above. The difference is that the three selection steps are now carried out with a hybrid objective implementing thermodynamic calculations. Inexpensive string criteria first define admissible candidates and sampling neighborhoods, while NUPACK-derived thermodynamic quantities bias and rank the sampled toehold sets, barcode sets, and toehold–barcode matchings. This approach retains the computational advantages of the string-based PyWinder generator, while allowing physically motivated equilibrium quantities to be considered. We will first define these thermodynamic metrics and then describe their implementation within the overarching generation algorithm.

Unless otherwise specified, thermodynamic generation uses the NUPACK 4.0.2.0 default DNA model with Sidewinder-specific salt settings $\mathcal{M} = (T = 50°C, [Na^+] = 0.154\,M, [Mg^{2+}] = 0.01\,M)$ and a working strand concentration $c = 10^{-8}\,M$. For two sequences $a$ and $b$, define $C(a, b)$ as the NUPACK expected equilibrium concentration of the duplex complex $(a, b)$ in a tube initialized with $[a] = [b] = c$ and with all complexes up to size two (i.e. the number of individual strands in a complex) considered in the partition set given the aforementioned settings. For candidate sequences $x$ and $y$, we define the thermodynamic off-target probability as:

$$P_{off}(x, y) = \frac{1}{c} \max\{C(x, \tilde{y}), C(\tilde{x}, y)\}.$$

Thus only the two sense–antisense orientations most directly analogous to misligation-prone Sidewinder interactions are considered at this stage; same-orientation duplexes and larger complexes are not used in candidate-stage scoring. For barcode candidates only, PyWinder will also compute the normalized on-target binding probability: $P_{on}(x) = C(x, \tilde{x})/c$.

The generator also computes NUPACK ensemble-defect quantities. The ensemble defect is the average number of incorrectly paired nucleotides at equilibrium compared to a target secondary structure. Let def$(X; \sigma)$ denote the ensemble defect of a strand complex $X$ relative to target structure $\sigma$. Moreover, let $\sigma =$ **duplex** denote a fully paired duplex structure (appropriate if $X$ is a complex composed of 2 strands of the same length) and $\sigma =$ **ss** denote a fully single-stranded structure with no secondary structure (appropriate if $X$ is a single stranded complex).

For a candidate sequence $x$, the on-target defect metric is:

$$D_{on}(x) = \max\{\text{def}((x,\tilde{x}); \textbf{duplex}), \text{def}((x); \textbf{ss}), \text{def}(\widetilde{(x)}; \textbf{ss})\},$$

so lower values indicate lower defect relative to the intended duplex state and the isolated single-stranded states.

For a pair of distinct candidate sequences $x$ and $y$, the off-target defect is:

$$D_{off}(x,y) = \min\{\text{def}(x,\tilde{y}; \textbf{duplex}), \text{def}(\tilde{x}, y; \textbf{duplex})\}.$$

Here larger values are preferred, because a large ensemble defect relative to a fully paired off-target duplex implies weaker support for that off-target bound state.

Thermodynamic toehold selection begins by building the same weighted edit-distance matrix $d_{toe}$ used by the string-based algorithm. For each candidate toehold vertex $x$, the generator calculates $D_{on}(x)$; for each candidate pair $x, y$ it calculates $P_{off}(x,y)$ and $D_{off}(x,y)$. These quantities define vertex and edge weights for biased path sampling. The initial toehold in a sampled path is drawn from its randomly chosen locus with vertex weight

$$W_v(x) = \exp[-D_{on}(x)].$$

Subsequent loci are still visited in random order, but candidates in the next locus are sampled with weights proportional to one plus the sum of edge weights to the already selected path vertices, where

$$W_e(x,y) = \exp\{\widehat{d}_{toe}(x,y) - 2P_{off}(x,y) + D_{off}(x,y)\}$$

and $\widehat{d}_{toe}$ is the normalized weighted edit-distance score. Thus thermodynamic toehold sampling favors candidates that are string-dissimilar, have low predicted off-target binding probability, have low on-target ensemble defect, and have high off-target ensemble defect.

After $K$ thermodynamically biased toehold paths have been sampled ($K = 1000$ by default), each path $P$ is scored by the tuple

$$S_{toe}^{thermo}(P) = (\max P_{off}, \overline{P}_{off}, \max D_{on}, \overline{D}_{on}, \min D_{off}, \overline{D}_{off}, \min d_{toe}, \overline{d}_{toe}).$$

The first four components are minimized and the final four are maximized by the same weighted rank-aggregation procedure described for string-based generation. The default rank weights are $2.0$ and $1.0$ for the maximum and average off-target probabilities, $1.0$ and $0.5$ for the maximum and average on-target defects, $1.0$ and $0.5$ for the minimum and average off-target defects, and $1.0$ and $0.5$ for the minimum and average weighted edit distances. This gives strongest priority to suppressing the worst predicted off-target concentration, while still preserving the string-orthogonality criteria that prevent the thermodynamic objective from dominating the search.

Given the selected thermodynamic toehold set $\mathcal{T}$, the barcode candidate pool is generated by the same SSM procedure used in string-based PyWinder generation. Barcode candidates are then evaluated with conventional edit distance, $P_{on}$, $P_{off}$, $D_{on}$, and $D_{off}$. The biased barcode sampler uses vertex weights

$$W_v(\beta) = \exp[-2D_{on}(\beta) + 0.5P_{on}(\beta)]$$

and edge weights

$$W_e(\beta_i, \beta_j) = \exp\{\widehat{d}_{bar}(\beta_i, \beta_j) - 2P_{off}(\beta_i, \beta_j) + 0.5D_{off}(\beta_i, \beta_j)\}.$$

Each sampled barcode set $B$ is ranked by

$$S_{bar}^{thermo}(B) = (\min P_{on}, \overline{P}_{on}, \max P_{off}, \overline{P}_{off}, \max D_{on}, \overline{D}_{on}, \min D_{off}, \overline{D}_{off}, \min d_{bar}, \overline{d}_{bar}).$$

The on-target probabilities, off-target defects, and edit distances are maximized; the off-target probabilities and on-target defects are minimized. Default rank weights are 0.5, 0.25, 2.0, 1.0, 2.0, 1.0, 0.5, 0.25, 1.0, and 0.5 in the order listed above.

The final thermodynamic selection step assigns the picked barcodes to the picked toeholds. For every candidate concatenation $z_{ij} = \tau_i \bullet \beta_j$, PyWinder computes the ensemble defect $D_{pair}(\tau_i, \beta_j)$ of the single strand $z_{ij}$ relative to a fully unpaired structure (**ss**). Pairings are sampled as bijections between $\mathcal{T}$ and $\mathcal{B}$, with the choice of a remaining toehold for a given barcode biased by $\exp[-D_{pair}(\tau_i, \beta_j)]$. Each complete bijection $\pi$ induces $z_i = \tau_i \bullet \beta_{\pi(i)}$ and is scored by

$$S_{pair}^{thermo}(\pi) = (\text{LCS}(\{z_i\}), \max_i D_{pair}(\tau_i, \beta_{\pi(i)}), \overline{D}_{pair}),$$

where LCS here is the longest common substring length among the concatenated toehold–barcode sequences. All three pairing metrics are minimized, with default weights $1.0$, $1.0$, and $0.5$, respectively. Once the thermodynamic toehold set, barcode set, and pairing have been selected, final Sidewinder strands are constructed exactly as in the string-based generator.

### String-based Sidewinder strand generation with thermodynamic filtering

The thermodynamic generator described above injects NUPACK calculations into each selection stage which significantly increases the computational cost of the algorithm as we perform thermodynamic calculations across each possible candidate toehold and barcode (**Fig. 1c**). PyWinder also supports a cheaper filtering mode in which the complete string-based pipeline is run repeatedly, and only these finished candidate libraries are evaluated and compared thermodynamically. This mode is useful when many fast string-generated designs can be produced, but would be too expensive to use thermodynamic scoring throughout the full combinatorial search in generation.

In this filtering mode, the string-based PyWinder generator is first run $M$ independent times to produce candidate Sidewinder libraries $\mathcal{L}_1, \ldots, \mathcal{L}_M$ ($M = 10$ by default unless otherwise specified). Each candidate library is saved, parsed, and checked to verify that the top strands reconstruct the input TSs and that the bottom strands reconstruct the corresponding reverse complements. For each candidate library ($\mathcal{L}_m$), the filter extracts the selected toehold–barcode concatenations

$$Z^{(m)} = z_1^{(m)}, \ldots, z_{n_m}^{(m)}, \qquad z_i^{(m)} = \tau_i^{(m)} \bullet \beta_i^{(m)},$$

and evaluates a library-specific full duplex off-target probability matrix $\widetilde{P}^{(m)}$ using the same default NUPACK model and concentration defined previously in the thermodynamic algorithm section.

For two distinct concatenated sequences $z_i^{(m)}$ and $z_j^{(m)}$ from $\mathcal{L}_m$, the filtering matrix uses the general duplex off-target probability

$$\widetilde{P}_{ij}^{(m)} = \frac{1}{c}\max\{C(z_i^{(m)}, z_j^{(m)}), C(z_i^{(m)}, \widetilde{z_j^{(m)}}), C(\widetilde{z_i^{(m)}}, z_j^{(m)}), C(\widetilde{z_i^{(m)}}, \widetilde{z_j^{(m)}})\}.$$

For diagonal entries, define $C_{2c}(a, a)$ as the expected concentration of the homodimer $(a, a)$ in a tube initialized with $[a] = 2c$. Then

$$\widetilde{P}_{ii}^{(m)} = \frac{1}{c}\max C_{2c}(z_i^{(m)}, z_i^{(m)}), C_{2c}(\widetilde{z_i^{(m)}}, \widetilde{z_i^{(m)}})$$

Thus, for each design, the filter checks for the predicted probability of an unintended duplex, including sense-sense, sense-antisense, antisense-sense, antisense-antisense, and self-dimer interactions. This is a deliberately conservative metric.

Each candidate library ($\mathcal{L}_m$) is assigned the scalar filtering score:

$$\Omega(\mathcal{L}_m) = \max_{1 \le i,j \le n_m} \widetilde{P}_{ij}^{(m)}$$

Libraries are ranked in increasing order of $\Omega$, such that a lower score indicates that the worst predicted toehold–barcode off-target interaction is smaller.

For the data presented in this manuscript, only the Foldtuned GFP pools used end-point thermodynamic filtering with $M = 10$ for the $10\%$ identity pool and $M = 25$ for the $20\%, 30\%, 55\%, 65\%$, and $70\%$ pools.

### Sidewinder strand generation speed comparison

A runtime comparison between the original NUPACK multitube design job method (*9*) and all PyWinder variants is shown in **Fig. 1c**. This comparison used $b = 18, t = 10$, and a max final oligo length of 165 bp. Designs with increasing numbers of barcode-toehold pairs were generated using increasing numbers of TS inputs drawn from the 24 secreted human protein coding sequences discussed. Timing was completed on a Linux workstation running Fedora 42 with kernel 6.18.5-100.fc42.x86_64, equipped with an AMD Ryzen 9 9950X3D CPU (16 physical cores / 32 hardware threads), and 128 GB DDR5 RAM configured at 3600 MT/s. PyWinder was run with 32 worker processes; NUPACK 4.0.2.0 was used and permitted to use a maximal number of threads on the machine.

### Oligo Purchasing

All assembly oligos were purchased from Integrated DNA Technologies (IDT) as oPools Oligo Pools at the 10 pmol/oligo synthesis scale. Assemblies have been successful using as low as 1 fmol input per oligo but the 10 pmol scale allowed for multiple experiments to be conducted using the same pool. PCR amplification primers under 60 bases were ordered from Integrated DNA Technologies with standard desalt purity and PCR amplification primers 60 to 120 bases were ordered from Millipore-Sigma with standard desalt purity. All oligos were shipped dry. All oligos are listed in **Supplementary Table 1**.

### Oligo phosphorylation and annealing

After centrifugation, oligo pools were resuspended in 100 µL of 1× T4 Ligase Buffer from New England Biolabs (NEB) and gently vortexed. An aliquot of the oligo pool was taken for phosphorylation in a 10 µL reaction of 1× T4 ligase buffer. By manufacturer's recommendation, 1 µL of T4PNK (NEB) phosphorylates 400 pmol of 5' DNA end. The volume of stock pool required for 400 pmol of oligo was phosphorylated with 1.1 µL of T4PNK in a 10 µL reaction at 37°C for 1 hour on a thermocycler.

Immediately after phosphorylation, the thermocycler temperature was raised to 98°C for enzyme deactivation and oligo denaturation. The temperature was then decreased at -1°C per minute to 25°C and held at 25°C or 4°C until use.

### Sidewinder Assembly

Assembly reactions were conducted in 50 µL final volume of 1× Hifi Taq Ligase buffer (NEB). The entire 10 µL phosphorylated, annealed oligo pool was added to 5 µL of 10× Hifi Taq Buffer and 33 µL of $H_2O$. The 48 µL mixture was placed on the thermocycler and 2 µL of Taq ligase (NEB) was added after the reaction increases to temperature directly on the thermocycler.

Three different assembly protocols are used in the manuscript where described.

#### Assembly protocol 1

1. 68°C for 5 min
    a. Add ligase
2. Drop temperature from 68°C to 37°C at -1°C per minute
3. Hold at 37°C for 20 minutes
4. Raise temperature back to 68°C
5. Repeat steps 2-4 for total of 4 cycles
6. Hold at 37°C for 8 hours.

#### Assembly protocol 2

1. 85°C for 5 min
    a. Add ligase
2. Drop temperature from 85°C to 37°C at -1°C per minute
3. Hold at 37°C for 20 minutes
4. Raise temperature back to 85°C
5. Repeat steps 2-4 for total of 4 cycles

#### Assembly protocol 3

1. 68°C for 5 min
    a. Add ligase
2. Drop temperature from 68°C to 37°C at -1°C per minute
3. Hold at 37°C for 20 minutes
4. Raise temperature back to 68°C

5. Repeat steps 2-4 for total of 10 cycles
6. Hold at 37°C for 8 hours.

### PCR amplification and purification

After assembly, repliQa HiFi ToughMix (Quantabio) was used for amplification of assemblies. We observed 1 $\mu$L of unpurified 3WJ assembly from the previous assembly step was sufficient template in a 50 $\mu$L PCR reaction for robust amplification. PCR reaction conditions were established according to manufacturer recommendations and predicted Tm of primers on SnapGene.

#### 24 construct specific amplifications

1. 98°C for 45 s
2. 98°C for 15 s
3. 68°C for 6 s
    a. Go to Step 2 for 40 cycles
4. 68°C for 5 min

#### Universal amplifications

1. 98°C for 45 s
2. 98°C for 15 s
3. 63°C for 5 s
4. 68°C for 5 s
    a. Go to Step 2 for 20-40 cycles
5. 68°C for 5 min

#### Evo 12.5 kb assembly amplicon

1. 98°C for 45 s
2. 98°C for 15 s
3. 64°C for 5 s
4. 68°C for 125 s
    a. Go to Step 2 for 40 cycles
5. 68°C for 5 min

Post PCR amplification, Gel extraction was done on assembled products which results in a purer product for downstream sequencing or cloning (**Fig. S2**). The Monarch DNA Gel Extraction Kit (NEB) was used according to the manufacturer's protocol for all samples prior to sequencing, except the explicitly stated Assembly 1 with standard purification. For this condition, purification of the PCR reaction was done using a QIAquick PCR Purification Kit (Qiagen).

### Hierarchical assembly

After verification that the universal amplicon contained each of the 20 target constructs, the single band was gel extracted and assembled via Golden Gate. The 20 fragments were assembled in a 50 µL reaction at approximately 8 nM (assuming equivalent concentration per construct) in 1× T4 DNA ligase buffer. The sample was placed at 37°C for 16 hr with 1 µL BbsI-HF (NEB) and 2 µL T4 Ligase (NEB). Post assembly, 1 µL of the reaction mixture was used as the template in a 50 µL PCR. Amplification was achieved across as many as 10 Golden Gate fragments.

More reliable amplification from the Golden Gate assembly was observed when amplifying the reaction in four 3 kb fragments so the decision was made to process these 4 amplicons for another round of *in vitro* hierarchical assembly. This second assembly step also allowed for easy addition of a fifth fragment containing an antibiotic resistance marker to be amplified and added to the 12.5 kb synthetic construct for more efficient integration into the genome. The 5 fragments were amplified with deoxyuracil containing primers and processed via USER cloning by first digesting at 37°C in a 32.22 µL reaction in 1× Cutsmart with 1 µL USER enzyme cocktail (NEB). The digests were purified via gel extraction and assembled at 8 nM in a 25 µL reaction of 1× Hifi Taq ligase buffer at 50°C for 16 hr with 1 µL Taq ligase. Post assembly, 1 µL of the reaction mixture was used as the template in a 50 µL PCR.

### DNA gel imaging

Gel electrophoresis was conducted with 1-2% agarose gels stained with Sybr Safe (Invitrogen, Thermo Fisher Scientific) in 0.5× TBE buffer (Genesee Scientific) run for 25 minutes at 135v. Main figure gels show 50 ng of template loaded measured using the Qubit 1x dsDNA High Sensitivity Assay Kit (Invitrogen, Thermo Fisher Scientific).

### 12.5kb sequence generation with Evo 2

A 3kb segment of *Escherichia coli* MDS42 genome was used as the prompt to initiate the generation of 20 sequences of 15 kb in length. We used the Evo 2 40B model with temperature of 0.7 and top-k of 4. With the same settings, 20 sequences of the same region of 15 kb were generated from the reverse direction also using a 3kb prompt. We compared the results with the reference MDS42 genome and chose the AI designed sequence with the highest identity in both directions and combined them into a 12.5kb targeted sequence for construction. To ensure the design was compatible with downstream processing, all BbsI restriction sites were replaced with silent mutations.

### Design of GFP Sequences

Foldtuning of ProtGPT2 was performed essentially as previously described with modifications as follows (*24, 31*). Experimental target structures of natural GFP-like proteins (n=480) and natural GFP-like sequences for preliminary evotuning (n=325, reduced to n=205 after clustering at 100% similarity) were obtained using Foldseek in fast TM-align mode to query reference structure pdb:1ema against the pre-computed PDB and UniRef50 databases respectively, filtering to structures with minimum 80% query coverage and alignment TM-score > 0.5. 2000 sequences were generated from ProtGPT2 for the evotuning round and for each of four rounds of foldtuning, resulting in 1530 *in silico*-validated candidates after structure prediction with 5.6-18.5% sequence identity to W.T. GFP.

Inverse-folding redesign of GFP was performed with ProteinMPNN (*32*), taking pdb:1ema as the input template, setting training backbone noise to 0.02 Å and sampling temperature to 0.3, and setting inference-time backbone noise to 0.0, 0.02, 0.05, 0.1, 0.15, 0.2, or 0.3 Å, generating 1000 sequences per parameter set, for 7000 sequences in total. Chromophore region amino-acid identities were restricted to S or T at position 65 (with equal probability), Y at position 66, and G at position 67; additional constraints were enforced at chromophore-maturation-influencing positions 96 (R with 1.0 probability) and 222 (E with 1.0 probability). Sequences were binned into quintiles by sequence identity to W.T. GFP, with quintile medians of 21.6%, 30.9%, 57.0%, 65.6%, and 70.0% identity giving the pool demarcations (20%, 30%, 55%, 65%, and 70%). Predicted structures for all 7000 variants were obtained from Boltz-2; within each quintile sequences were ranked according to the geometric mean of predicted structure pLDDT (model confidence score) and TM-score vs pdb:1ema (structural similarity to input template).

### Transformation and cloning

For the GFP libraries, the universal amplicon was amplified using deoxyuracil-containing (dU) primers with repliQa HiFi ToughMix. A plasmid backbone containing Tetracycline resistance gene, p15a origin, and T7 promoter was amplified also using dU primers. All primers are listed in **Supplementary table 1** and fasta reference sequences for the library targets and backbone is listed in **Supplementary table 2**.

The GFP library amplicons and backbone were digested individually in 50 μL reactions in 1× Cutsmart buffer with 1 μL USER (and 4 μL DPN1 for the backbone fragment) at 37°C for 1 hour and 80°C for 20 minutes. Digested samples are then purified with Monarch DNA Gel Extraction Kit. Assembly into the backbone is conducted at 6 nM backbone 18 nM insert in 1× Hifi Taq Buffer at 50°C for 16 hr with 2 μL Taq ligase. Assembly is then purified with QIAquick PCR Purification Kit (Qiagen) and eluted in 30 μL H2O.

10 μL of the purified sample is electroporated into 100 μL of electrocompetent DH10B cells. Transformation is then recovered for 1 hr in 2 mL of Luria–Bertani (LB) media. After recovery, the 2 mL of recovery was inoculated into 48 mL LB with 10 μg/mL tetracycline and grown to approximately OD 1.0. Five mL of the sample was then spun down and miniprepped and PCR'd with the universal amplification primers for sequencing to get the no induction sequence distribution.

This miniprep was used to transform the library into electrocompetent DH10B containing a genomically integrated T7 polymerase, recovered for 2 hr, and plated on 100 μM IPTG, 10 μg/mL Tetracycline, LB agar plates at 1:100 dilution and grown overnight at 37°C . These plates were then scraped with 4 mL LB, miniprepped, and PCR'd with the universal amplification primers for sequencing to get the induction sequence distribution.

### Sequencing analysis

PacBio HiFi sequencing (GENEWIZ) was used to obtain high-confidence, full-molecule long reads for all samples. Raw HiFi reads in FASTQ format were processed through a custom bioinformatics pipeline performing three analyses: single-nucleotide polymorphism (SNP) profiling, fragment classification, and junction analysis.

For SNP profiling: reads were aligned to reference sequences using the Smith-Waterman local alignment algorithm (33) with a match score of 5, mismatch penalty of 4, gap open penalty of 10, and gap extend penalty of 0.5. Only reads with a mean Phred quality score of greater than or equal to Q39.5 were retained. At the base level, only positions with a Phred score of greater than or equal to Q40 were included to minimize errors due to sequencing. Reads were further filtered to require greater than or equal to 90% alignment identity and greater than or equal to 80% aligned fraction of the reference length. The SNP rate at each position was calculated as the total number of non-reference events (substitutions, deletions, and insertions) divided by total coverage at that position. Homopolymeric runs had the total insertions and deletions distributed evenly across the stretch of the run to account for discrepancies in assigning insertions and deletions to one of a particular position in a homopolymeric run. Positions with no observed errors were plotted in the distribution with error rate $10^{-7}$. The raw number of mutation free molecules observed for each construct is reported in **Source Data**. The calculations do not consider the terminal 50 bases on either end of the construct due to noise at the end of the molecule during sequencing. The rate of nucleotide perfect clones reported in the results is calculated as the ratio between the number of mutation-free molecules observed and the average coverage of each base for that construct.

Fragment-level analysis: read sequences were aligned to all fragment references using BLASTn (*4*). Reads returning no hits to any fragment were classified as unusable. A complete assembly was assigned when all fragments were detected in the correct order of the gene sequence; assemblies with correct connections between fragments but missing a subset of fragments with no misconnections were classified as partial. Reads containing fragments joined in an incorrect order were classified as "other" including PCR and sequencing artifacts or incorrect assemblies. This is the same classification scheme used in our prior publication (9). Fragment hits were required to cover at least 80% of the fragment reference length. For pooled samples containing multiple reference constructs, reads were additionally assessed for cross-reference crosstalk.

Junction analysis: a junction was defined as 25 base pairs flanking both the 3' and 5' ends of each Sidewinder junction. For each sample, all possible junctions, including both correct ligations and potential mis-ligations, were generated from the fragment references and aligned to the raw reads using BLASTn with sensitive parameters. Junction hits were filtered using query coverage (greater than or equal to 80%) and percent identity (greater than or equal to 80%) thresholds. For reads identified as potential mis-ligations, they were counted as true mis-ligations if a read contained a seamless junction at the exact 6 base pairs on either side of the ligation point between non-partnered fragments. This helped to remove hits on non-junction regions of the constructed sequences which happened to share 80% similarity to a possible junction combination.

For the universal amplification of the 20 Evo designed sequences, in some instances an intended 3WJ junction would be present in the universal buffer region which was partially shared by all 20 sequences resulting in a similar false identification of non-junction regions as mis-ligations. As a result, the reported junction misconnection rates in **Figure 5g**, and **Figure 5k** exclude 3' and/or 5' ends that are partially or fully composed in the universal buffer region. Further, for all reported junction misconnection rates, the 3' of the coding oligo of the first fragment of a construct and the 5' end of the last fragment of a construct were not counted as misconnections because these are blunt ends and not 3WJs.

### Statistics and reproducibility

Bray-Custis distance matrices for principal coordinate analysis (PCoA), permutational multivariate analysis of variance (PERMANOVA), and permutational analysis of multivariate dispersions (PERMDISP) were calculated from normalized target read proportions. PCoA was performed using the PCoA function in scikit-bio. PERMANOVA and PERMDISP were performed using the permanova and permdisp functions in scikit-bio, respectively, each with 9,999 permutations.

### Supplementary Figures

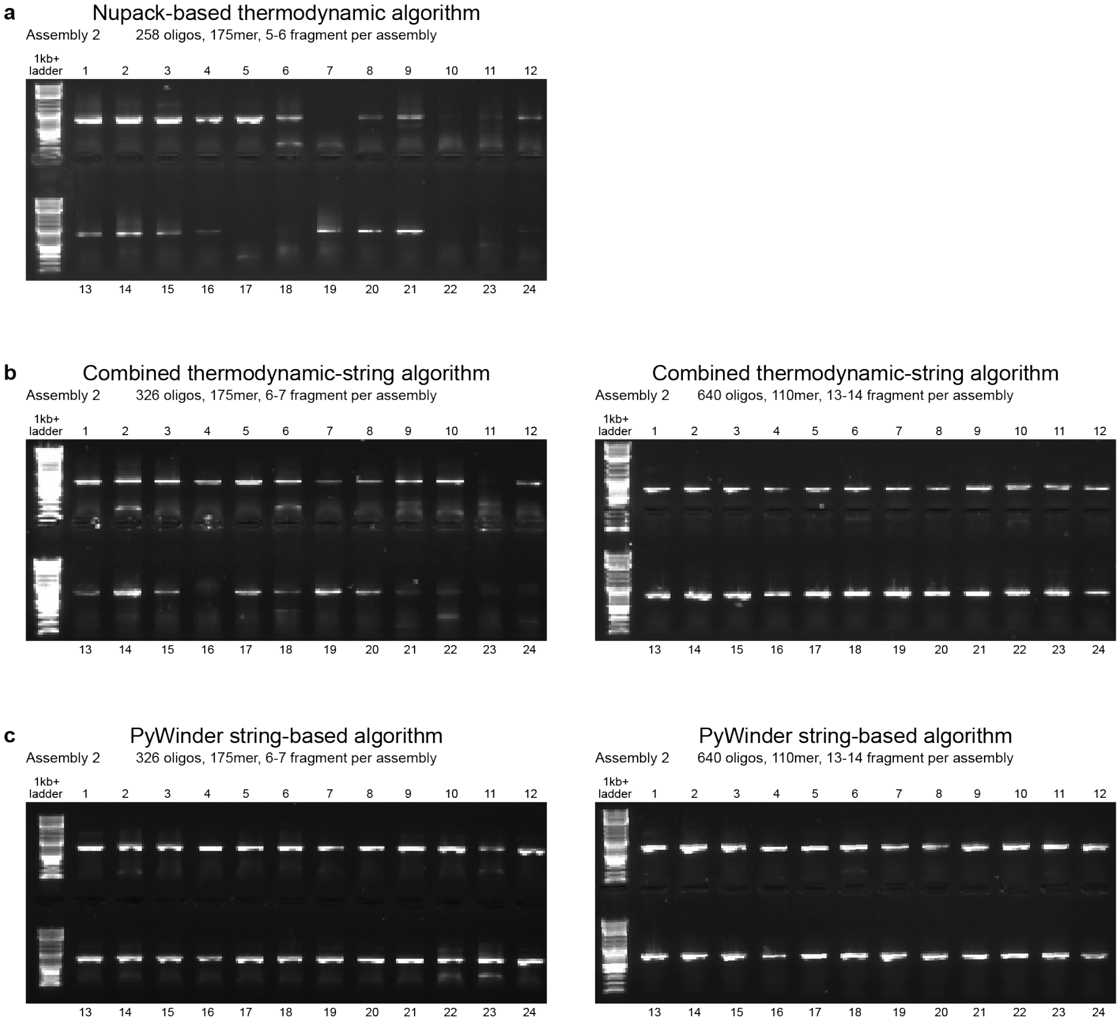


**Fig. S1. Oligo length and barcode design algorithm influence assembly performance a)** DNA agarose gel depicting 1 µL of the final PCR products for each of 24 individual constructs when the component Sidewinder oligos were generated with the NUPACK-based thermodynamic algorithm. **b)** DNA agarose gel depicting 1 µL of the final PCR products for each of 24 individual constructs when the component Sidewinder oligos were generated with the combined thermodynamic-string based algorithm for 175mer oligos (left) and 110mer oligos (right). **c)** DNA agarose gel depicting 1 µL of the final PCR products for each of 24 individual constructs when the component Sidewinder oligos were generated with the PyWinder string-based algorithm for 175mer oligos (left) and 110mer oligos (right).

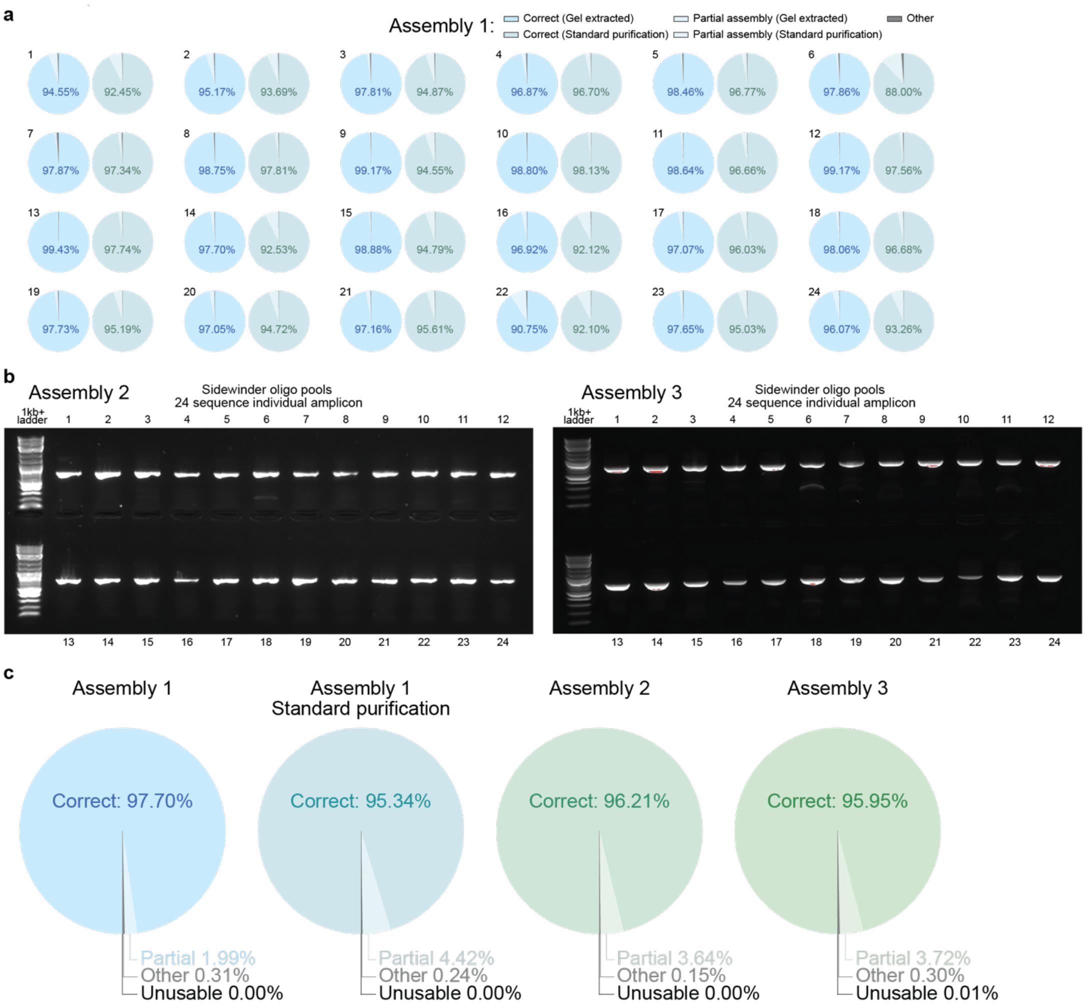


**Fig. S2. Sidewinder construction from oligo pools is reproducible across assembly protocols with a high proportion of correct assemblies**. Raw data available in **Source Data**. **a)** Fragment level PacBio sequencing analysis for each of the 24 individual constructs for Assembly 1 depicting correct assemblies (saturated), partially aligned products (a subset of fragments in the correct order) (unsaturated), and PCR artifacts, sequencing artifacts, or incorrect assemblies (grey). Gel extracted samples (blue) yield higher proportion of correctly assembled product compared to standard purification (teal) **b)** DNA agarose gel depicting 1 μL of the final PCR products for each of 24 individual constructs with a single strong target band for Assembly 2 and Assembly 3. **c)** Fragment level PacBio sequencing analysis combining all reads for all 24 individual amplicons for different assembly or purification conditions, depicting correct assemblies (saturated), partially aligned products (unsaturated), PCR artifacts, sequencing artifacts, or incorrect assemblies (grey), and unusable reads (not aligning to any reference) (black).

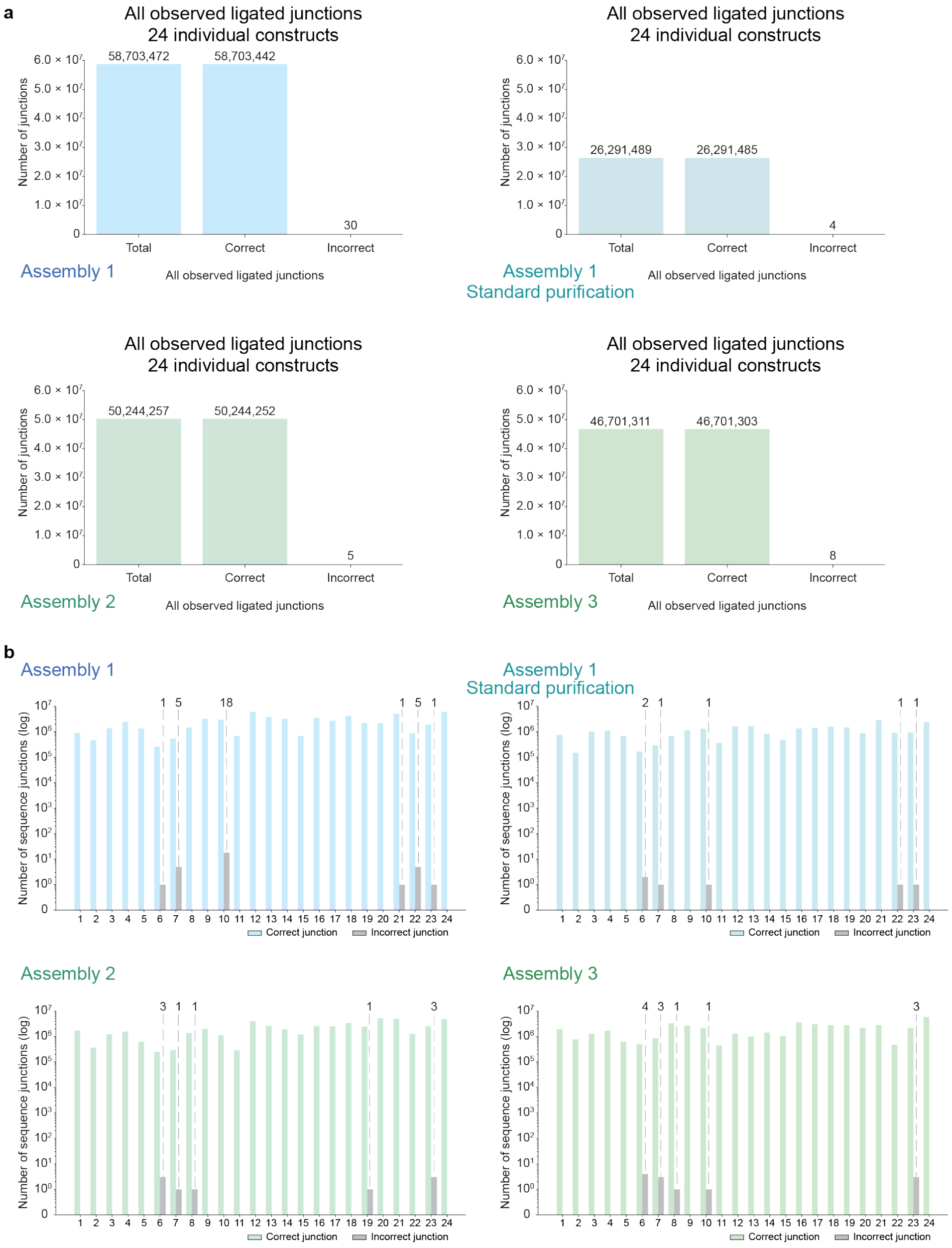


**Fig. S3. Sidewinder construction from oligo pools enables misconnection rates less than 1 in 10,000,000**. Raw data available in **Source Data**. **a)** Junction level PacBio sequencing analysis from unfiltered data depicting the global count of all observed junctions for all 24 construct specific amplicons for different assembly or purification conditions, depicting total junctions, correctly ligated junctions, and incorrectly ligated junctions. **b)** Junction level PacBio sequencing analysis from unfiltered data depicting all observed junctions for each of the 24 construct specific amplicons for different assembly or purification conditions the number of correctly ligated junctions (colored), and incorrectly ligated junctions (grey) for different assembly or purification conditions on log scale. Sequences with no observed mis-ligated junctions do not display a bar. Inter-construct mis-ligations are counted as misconnections for both constructs when visualized by individual sequence.

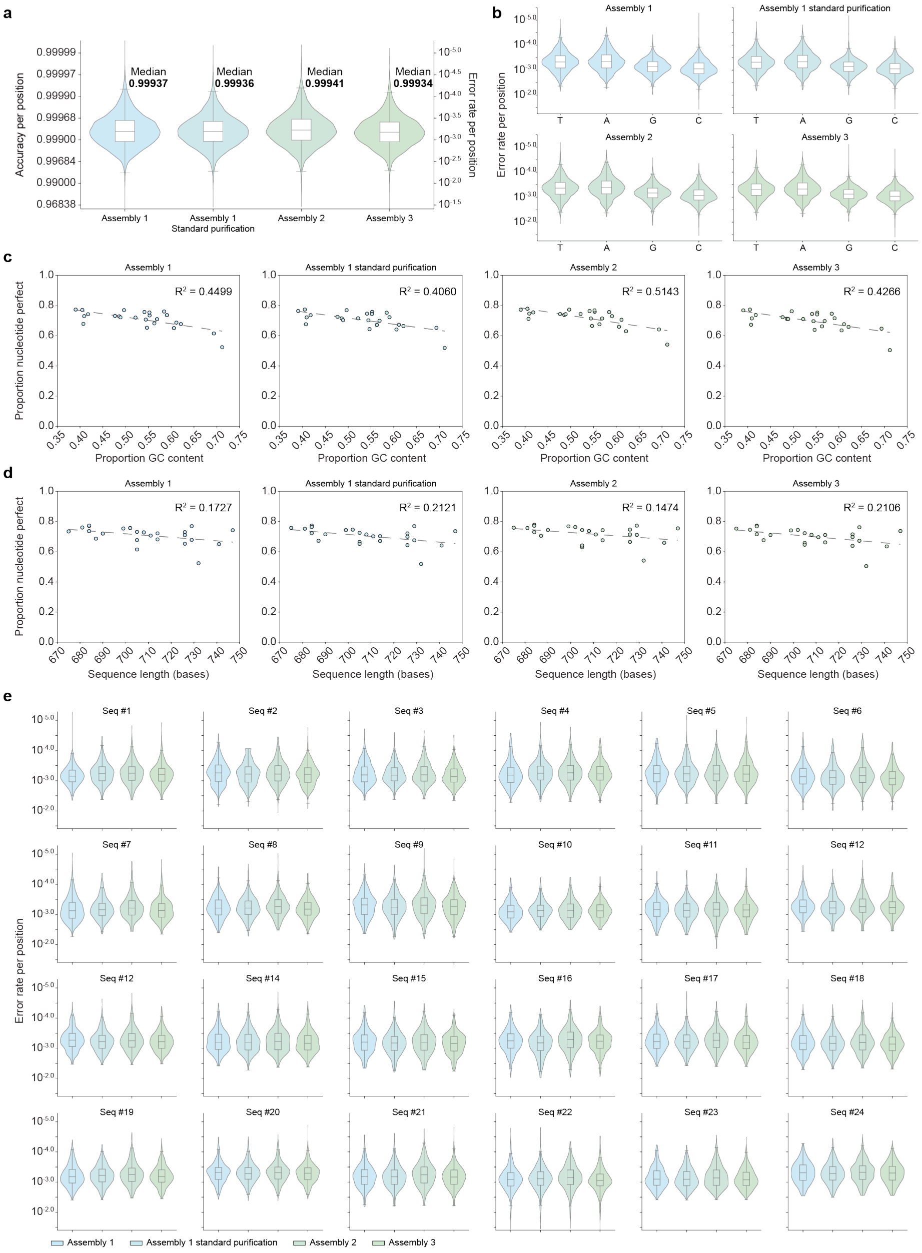


**Fig. S4. The per-position SNP frequency remains consistent across constructs and assembly protocols**. Summary statistics for all plots available in **Source Data**. **a)** Base level PacBio sequencing analysis depicting violin plots and nested box and whisker plots showing the distribution of per-base accuracies for all positions of all 24 constructs separated by assembly protocol. The median error rates are 1 in 1,587, 1,563, 1,695, and 1,515 for Assembly 1, Assembly 1 with no gel extraction, Assembly 2, and Assembly 3 respectively. **b)** Base level PacBio sequencing analysis depicting violin plots and nested box and whisker plots showing the distribution of per-base accuracies for all positions of all 24 constructs separated by assembly protocol and base identity. The base identity corresponds to the antisense strand relative to the reference sequence as the coding strand of the Sidewinder fragment is the template for the final 2WJ construct. **c)** Scatter plots plotting the relationship between proportion of SNP free reads assigned to each sequence versus GC content for all 24 constructs separated by assembly protocol. **d)** Scatter plots plotting the relationship between proportion of SNP free reads assigned to each sequence versus sequence length for all 24 constructs separated by assembly protocol. **e)** Base level PacBio sequencing analysis depicting violin plots and nested box and whisker plots showing the distribution of per-base accuracies for all positions of each of the 24 constructs separated by sequence and assembly protocol.

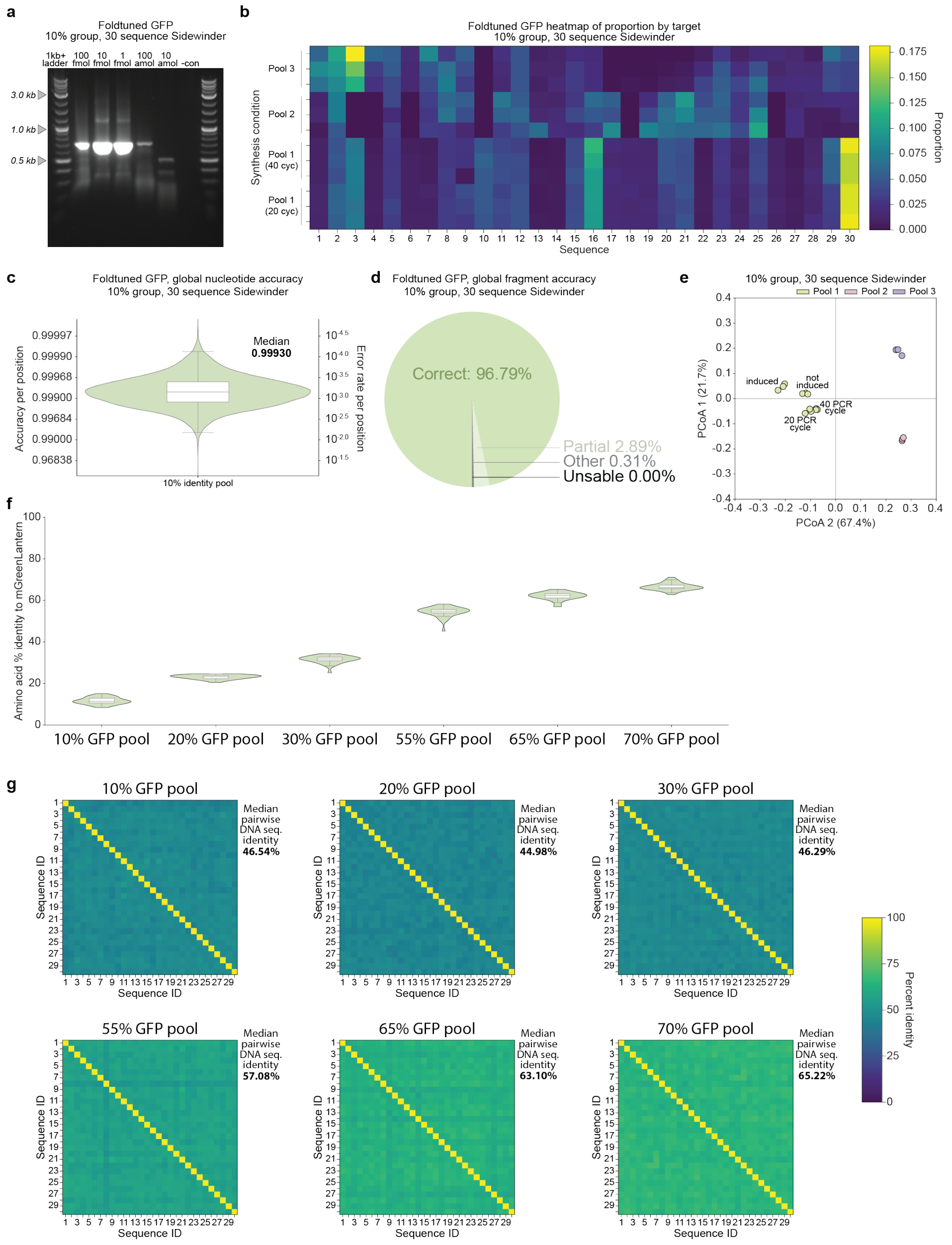


**Fig. S5. Sidewinder from oligo pools reliably assembles constructs in parallel at low concentrations, and with high inter-sequence identity**. Summary statistics available in **Source Data**. **a)** DNA agarose gel depicting 5 μL of the final PCR products for universal amplification at varying input oligo amounts. **b)** Heat map depicts the proportion of Oxford Nanopore sequencing reads assigned to each of the 30 sequences in the 10% identity pool universal amplicon after the 3WJ removal and PCR amplification with 20, 40 cycles, with 3 oligo synthesis replicates each done in triplicate. **c)** Base level PacBio sequencing analysis depicting a violin plot and nested box and whisker plot showing the global distribution of per-base accuracies for all positions of all 30 constructs in the 10% similarity group. **d)** Fragment level PacBio sequencing analysis combining all reads for all 30 sequences in the 10% group depicting correct assemblies (green), partially aligned products (a subset of fragments in the correct order) (light green), PCR artifacts, sequencing artifacts, or incorrect assemblies (grey), and unusable reads (not aligning to any reference) (black). **e)** Principal coordinate analysis (PCoA) plot based on Bray–Curtis distances for the 10% similarity group. Pool 1, 2, and 3 are three independently ordered and synthesized oligo pools of the exact same oligo sequences. Each pool was assembled in triplicate. Pool 1 was amplified via PCR with both 20 and 40 cycles and also transformed into cells both with and without induction. **f)** Violin plot and nested box and whisker plot showing the percent identity at the amino acid level of each of the 30 sequences to mGreenLantern GFP within each percent identity pool. **g)** Heatmap depicting pairwise DNA sequence alignment between each of the constructs within the 10%, 20%, 30%, 55%, 65%, and 70% similarity groups.

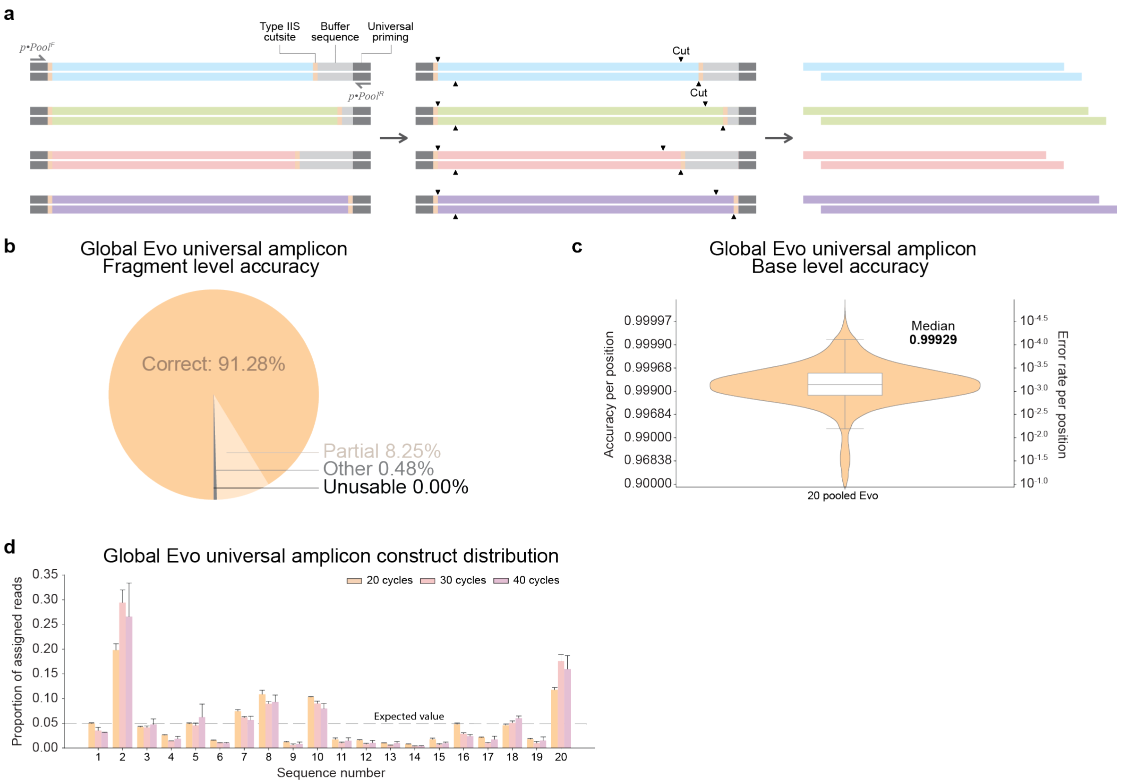


**Fig. S6. Constructs are designed for universal amplification with equivalent length and restriction sites for downstream processing, yielding highly accurate and robust assembly reactions. a)** The schematic depicts a universal priming region, external to a buffer sequence of variable length to match the longest constructed target. After amplification, a Type IIS restriction site is used to cut internally, removing all non-coding DNA, preparing all fragments for downstream processing simultaneously. **b)** Fragment level PacBio sequencing analysis combining all reads for all 20 sequences in the universal amplicon prior to hierarchical assembly depicting correct assemblies (orange), partially aligned products (a subset of fragments in the correct order) (light orange), PCR artifacts, sequencing artifacts, or incorrect assemblies (grey), and unusable reads (not aligning to any reference) (black). **c)** Base level PacBio sequencing analysis depicting a violin plot and nested box and whisker plot showing the global distribution of per-base accuracies for all positions of all 20 constructs prior to hierarchical assembly. **d)** Bar plot depicts the proportion of Oxford Nanopore sequencing reads assigned to each of the 20 sequences in the universal amplicon after the 3WJ removal and PCR amplification with 20, 30, and 40 cycles, each done in triplicate.

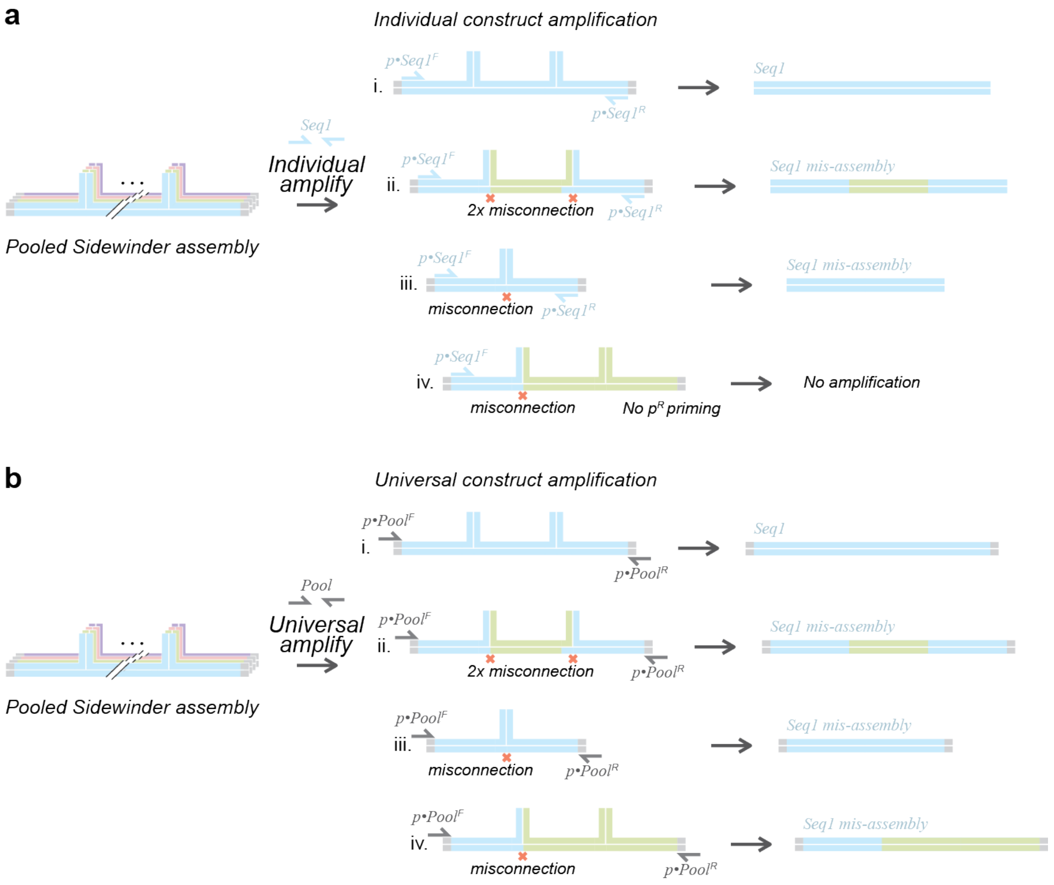


**Fig. S7. Hypothesized source for increase in measured misconnection rate when conducting universal construct amplification compared to individual construct amplification. a)** Schematic depicting construct specific amplification using primers which prime to just one of the sequences in a pool. Amplification requires the presence of both the first and last fragment of the sequence corresponding to the amplification primers. This is only the case when (**i**) the construct is correctly assembled, (**ii**) two independent misconnections occur within the same molecule, or (**iii**) a single misconnection occurs between fragments of the sequence corresponding to the amplification primers. **b)** Schematic depicting universal construct amplification using primers which prime to all the sequences in a pool. Amplification requires the presence of any first and last fragment of any construct in the pool. This is the case in the previous scenarios (**i-iii**) as well as scenario (**iv**) where a single misconnection occurs and is amplified by the universal amplification primers.
